# Lift observation conveys object weight distribution but partly enhances predictive lift planning

**DOI:** 10.1101/2020.06.21.163352

**Authors:** Guy Rens, Jean-Jacques Orban de Xivry, Marco Davare, Vonne van Polanen

**Author notes:** **Corresponding Author:** Guy Rens, The Brain and Mind Institute, University of Western Ontario, Ontario N6A 3K7, Canada. These authors contributed equally (Marco Davare, Vonne van Polanen). **Conflict of interest:** The authors declare to have no conflict of interest.

## Abstract

Observation of object lifting allows updating of internal object representations for object weight, in turn enabling accurate scaling of fingertip forces when lifting the same object. Here, we investigated whether lift observation also enables updating of internal representations for an object’s weight distribution. We asked participants to lift an inverted T-shaped manipulandum, of which the weight distribution could be changed, in turns with an actor. Participants were required to minimize object roll (i.e., ‘lift performance’) during lifting and were allowed to place their fingertips at self-chosen locations. The center of mass changed unpredictably every third to sixth trial performed by the actor and participants were informed that they would always lift the same weight distribution as the actor. Participants observed either erroneous (i.e., object rolling towards its heavy side) or skilled (i.e., minimized object roll) lifts. Lifting performance after observation was compared to lifts without prior observation and to lifts after active lifting, which provided haptic feedback about the weight distribution. Our results show that observing both skilled and erroneous lifts convey an object’s weight distribution similar to active lifting, resulting in altered digit positioning strategies. However, minimizing object roll on novel weight distributions was only improved after observing error lifts and not after observing skilled lifts. In sum, these findings suggest that although observing motor errors and skilled motor performance enables updating of digit positioning strategy, only observing error lifts enables changes in predictive motor control when lifting objects with unexpected weight distributions.

**New and noteworthy:** Individuals are able to extract an object’s size and weight by observing interactions with objects and subsequently integrate this information in their own motor repertoire. Here, we show that this ability extrapolates to weight distributions. Specifically, we highlighted that individuals can perceive an object’s weight distribution during lift observation but can only partially embody this information when planning their own actions.

## 1. Introduction

Skilled object manipulation not only relies on haptic feedback but also on anticipatory mechanisms (Johansson and Westling 1988; Johansson and Westling 1984). It has been argued that when individuals perform hand-object interactions, they form an ‘internal sensorimotor object representation’ which can then be retrieved to predictively plan fingertip forces for future object manipulations (Johansson and Westling, 1988; Gordon et al., 1991).

Much evidence has been given about the pivotal function of hand-object interactions for the formation of an internal object representation (Baugh et al. 2012; Fu et al. 2010; Gordon et al. 1991; Johansson and Westling 1984; Lukos et al. 2007). However, other studies have proposed that humans are able to generate similar representations when observing object lifting performed by others. For instance, previous studies have demonstrated that individuals are able to accurately estimate object weight during observed object lifting (Bingham, 1987; Runeson & Frykholm, 1981) and primarily rely on lift duration in doing so (Hamilton et al., 2007; Shim & Carlton, 1997). Furthermore, it has been demonstrated that this information can also be used to update the sensorimotor representation and improve predictive lift planning; After observing someone making typical lifting errors (e.g., using too much force when lifting a light object caused by overestimating its weight), lifting errors made by the second individual can be reduced (Meulenbroek et al., 2007) or even eradicated (Reichelt et al., 2013). However, both these studies only focused on the observation of salient movement errors and not on skilled lift performance.

To our knowledge, only a few studies compared how observing skilled or erroneous lifts mediate predictive object lifting. For instance, using the size-weight illusion, Buckingham et al. (2014) highlighted that predictive force scaling is improved after observing erroneous lifts compared to skilled ones: When participants had to lift a large, but unexpectedly light object for the first time, those who observed typical overestimation errors on the same object would make smaller lifting errors compared to those who observed skilled lifts. These findings and some of our previous results (Rens and Davare 2019) suggest that observing erroneous lifts conveys weight-related information better than skilled ones. In Rens and Davare (2019), participants were required to lift an object in turns with an actor. The object weight changed every n^th^ trial performed by the actor. Accordingly, participants could not predict the weight change but could have relied on observing the actor’s lifts to estimate object weight. Although our results showed that participants’ lift performance improved after observing skilled lifts, this improvement was smaller compared to when they observed lifting errors. These findings were supported by a later study of ours: Even though observation of skilled lifting enabled participants to improve their own lift planning, they were still biased by the object size during lift execution (Rens et al., 2020).

It is important to note that the studies mentioned above only considered object weight and size for investigating how lift observation improves predictive lift planning in the observer. Critically, when an object’s weight is asymmetrically distributed and digit positioning is not constrained (in contrast to previous studies such as Buckingham et al. 2014; Reichelt et al. 2013; Rens et al. 2020; Rens and Davare 2019), meaningful object interactions not only require fingertip forces to be anticipatorily scaled to the object weight but also to the external torque induced by the weight distribution. For instance, when carrying trays with multiple objects or when using tools with an asymmetrical weight distribution (e.g., a hammer), a valid digit force-coordination pattern is essential for skillful manipulation. Furthermore, it has been established that individuals often base their hand-object interaction on ‘end-state comfort’. That is, hand placement on objects is based upon the planned action after object contact (Cohen & Rosenbaum, 2004). As such, it has been argued that planning of hand-object interactions takes specific criteria, such as end-state comfort in account. Specifically, for objects with asymmetrical weight distributions, appropriate compensatory torque needs to be generated to offset the external torque and minimize object roll and to allow for skilled interactions.

Lukos et al. (2007) demonstrated that individuals can update their sensorimotor representation for an object’s weight distribution, in turn predictively generating an appropriate amount of compensatory torque. Fu et al. (2010) extended on their findings by showing that individuals appropriately scale their fingertip forces in function of digit positioning: When digit positioning is constrained, individuals are able to accurately scale their fingertip forces according to the fixed contact points. Conversely, when digit positioning is unconstrained, individuals are also able to accurately scale their fingertip forces in function of their self-chosen contact points. As such, these findings suggest that individuals can generate many equally valid digit position-force coordination patterns to minimize object roll during object lifting. Furthermore, successful force-positioning coordination patterns depend on the interaction between predictive force planning and sensory feedback about digit placement: Although digit positioning is planned predictively, individuals still demonstrate trial-to-trial variability (Fu et al., 2010). As a result, it is necessary to update the planned fingertip forces in function of this digit positioning variability.

With respect to end-state comfort, it is to note that all digit position-force coordination patterns might not be equal. For instance, Fu et al. (2010) showed that when lifting an object with asymmetrical weight distribution, positioning the fingertips furthers apart requires individuals to exert less force as the increased fingertip distance creates a ‘lever’ over which it is easier to generate compensatory torque. Accordingly, it is plausible that when individuals can position their fingertips without constraints, they would prefer a digit positioning strategy that requires less force to minimize object tilt. Taken together, this suggests that, when lifting asymmetrical weight distributions, individuals need to weigh the (dis)advantages of placing their contact points further apart or closer together as digit positioning and force scaling might contribute differently to end-state comfort.

To our knowledge, it has never been investigated whether lift observation can improve predictive lift planning for objects with an unknown weight distribution as previous studies (Buckingham et al. 2014; Meulenbroek et al. 2007; Reichelt et al. 2013; Rens and Davare 2019) only investigated the effect of lift observation on object weight-driven predictive lift planning. Importantly, many daily life hand-object interactions are performed in social settings (e.g., passing trays during dinner). When receiving an object either directly (hand-to-hand) or indirectly (placed within reach), the ‘receiver’ is unlikely to position their fingertips on the exact locations used by the ‘giver’. Accordingly, the receiver needs to estimate the object’s weight distribution based on observation and predictively plan a correct digit force-positioning coordination pattern to allow for skilled manipulations.

In the present study, we wanted to extend on the abovementioned literature by investigating (a) whether lift observation can convey critical information about an object’s weight distribution and (b) to which extent observing either lifting performance type (erroneous or skilled) mediates predictive lift planning better. For this, we asked participants to grasp and lift an inverted T-shaped manipulandum in turns with an actor. This object’s center of mass could be changed without the participants’ knowing. Participants were required to minimize object roll during lifting by generating an appropriate amount of compensatory torque. In addition, participants could place their fingertips at self-chosen locations requiring them to scale their fingertip forces in function of their self-chosen positioning. As such, participants could generate many equally valid digit position-force coordination patterns. The center of mass changed unpredictably every third to sixth trial performed by the actor, but participants were informed that they would always lift the same weight distribution as the actor. As such, participants could potentially estimate the object’s center of mass during observed object lifting, update their sensorimotor representation and subsequently plan their own lifting action correctly.

To investigate differences between performance types, we paired participants either with a naïve or informed actor. Naïve actors could not predict the center of mass change, making them unable to anticipatorily generate the appropriate amount of compensatory torque thus causing the inverted T-shape to roll towards its heavy side during lifting (‘erroneous’). In contrast, the informed actor could predict the center of mass change and subsequently generate enough compensatory torque for minimizing object roll during lifting (‘skilled’). As a result, the center of mass change was potentially indicated by either an erroneous (naïve actors) or skilled lift (informed actor). We hypothesized, in line with previous studies (Meulenbroek et al., 2007; Reichelt et al., 2013; Buckingham et al., 2014) that observation of lifting errors would improve predictive lift planning. In line with Rens and Davare (2019), we hypothesized that observation of skilled lifts would also improve predictive lift planning albeit in a smaller manner than observation of erroneous lifts.

## 2. Methods

### 2.1 participants

24 participants were recruited from the student body of KU Leuven (Belgium) and divided into two groups of 12 participants each. The first group (skilled observation group) consisted of 12 dyads in which each participant was paired with the experimenter (9 females and 3 males; mean age = 22.28 ± 0.65). The second group (error observation group) consisted of 6 dyads in which each participant was paired with another participant (9 females and 3 males; mean age = 21.38 ± 0.62). Participants in the second group did not know their paired partner in advance. All participants were right-handed (self-reported), had normal or corrected-to-normal vision, were free of neurological disorders and had no motor impairments of the right upper limb. The study was conducted in accordance with the declaration of Helsinki and was approved by the local ethical committee of the Faculty of Biomedical Sciences, KU Leuven. Participants gave written informed consent and were financially compensated (20 euro). We excluded the data of one dyad (both actor and participant data) in the skilled observation group due to errors in data. The force/torque sensors were incorrectly attached to the OEM board (see below), causing the data of the second session to be corrupted. The participant (female) declined to redo the second session. Because of this, we decided to also exclude the collected data of the first session due to the limited number of trials. As such, we had 12 participants in the error observation group and 11 in the skilled observation group.

### 2.2 data acquisition

For the present study, we used a custom-built carbon fiber ‘inverted T-shape’ grip-lift manipulandum (Arsalis, Belgium; for all object dimensions see Figure 1). The manipulandum consisted of a horizontal basis and a vertical block to which two 3D force/torque (F/T) sensors were attached. On each force sensor a cover plate (height × width: 140 × 53 mm) with a center protruding surface (height × width: 140 × 20 mm) was mounted to block the view of the F/T sensors. Both protruding surfaces were covered with fine-grained sandpaper (p600) and the horizontal distance between them was 48mm. During the experiment, participants and experimenter were allowed to place their thumb and index finger freely (precision grip) but only on the protruding surfaces (i.e., ‘the graspable surfaces’). The horizontal basis of the manipulandum consisted of three compartments which enabled the placement of 3D-printed cuboids that were identical in appearance (height x width x depth: 55 x 35 x 40 mm).

**Figure 1.**
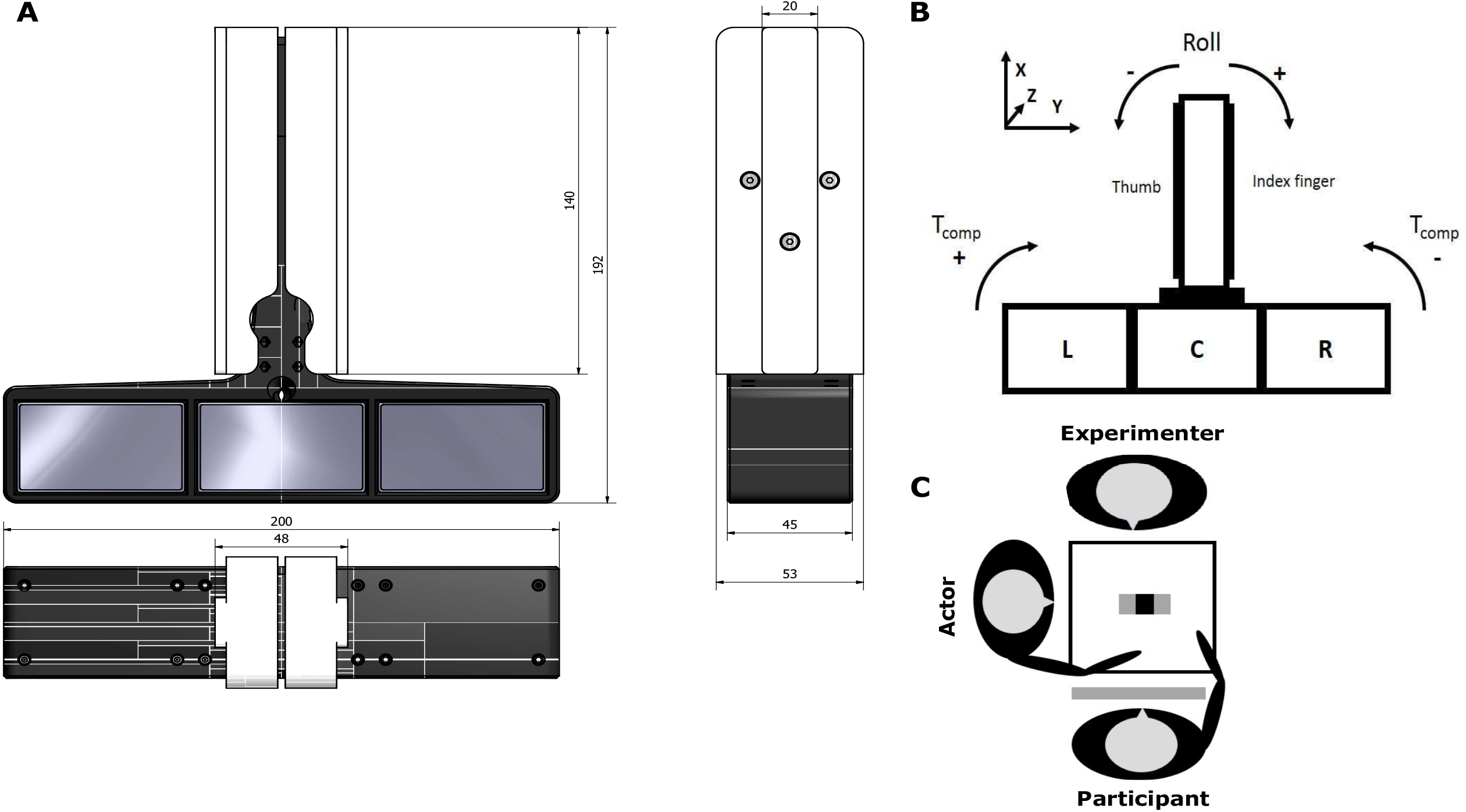
Manipulandum and set-up. **A**. Frontal, side and top down view of the ‘inverted T-shape’ manipulandum with dimensions (in mm). The manipulandum consisted of a vertical (white) component and a horizontal base (black). The horizontal base consisted of 3 compartments in which the 3D printed cuboids could be placed. **B**. Schematic drawing of the manipulandum with the three compartments indicated with L, C and R standing for left, center and right, respectively. Counterclockwise and clockwise roll were defined as negative and positive, respectively. Compensatory torque, i.e., torque generated by participants to offset object roll was defined as the inverse of object roll, i.e., positive and negative for clockwise and counterclockwise compensatory torque, respectively. X, Y and Z indicate the frame of reference for the force/torque sensors in the vertical component. **C**. The participant and actor (Skilled observation group: informed actor; Error observation group: naïve actor) were seated next to each other at a table on which the manipulandum was positioned and a switchable screen was placed in front of the participant’s face.

The manipulandum, without cuboids, weighed 3.95 N. Two cuboids were hollow and weighed 0.24 N each, the third one was filled with lead particles and weighed 4.24 N. The total weight of manipulandum and cuboids was 8.67 N. Inserting the heavy cuboid in the left, center or right compartment would induce an external torque of −245, 0 and +245 Nmm, respectively. Prior to the start of the study, the external torque was calculated in the following manner: For each center of mass, the manipulandum was held in the air with its base as horizontally as possible for 3 seconds. This was repeated 10 times (using different digit placement each repetition for when the heavy cuboid was in the side compartments). The amount of compensatory torque, i.e., amount of torque to keep the base horizontal, was calculated as the grand mean of the means of all repetitions for each center of mass location separately. Last, the external torque was defined as the opposite of the compensatory torque (e.g., compensatory torque for left center of mass = 245 Nmm, accordingly the external torque for the same center of mass = −245 Nmm).

In the present study we used two ATI mini-40 SI-40-2 F/T sensors (force range: 40, 40 and 120 N for x-, y- and z-axes respectively; force resolution: 0.01 N; torque range: 2 Nmm; torque resolution: 0.0005 Nmm) (ATI Industrial Automation, USA). In addition, a ± 3 g three-dimensional accelerometer (ADXL335, Analog Devices, USA) was mounted on top of the vertical block and was hidden beneath the cover plates. Both F/T sensors and the accelerometer were connected to the same NI-USB 6221 OEM board (National Instruments, USA) which was connected to a personal computer. Data was acquired using a custom-written MATLAB script (Mathworks, USA) and sampled at 1Khz. Last, F/T sensors and the accelerometer were calibrated by the developer (Arsalis) in accordance with the applicable QTI procedures. Briefly, F/T sensors and the accelerometer were attached to the manipulandum. In this condition, F/T sensors were loaded with calibrated weights to quantify their accuracy in stationary conditions. Similarly, the manipulandum was kept stationary in multiple orientations to quantify the accelerometer’s accuracy in three-dimensional space. The maximum amount of reported error for the force, torque and accelerometer components were 1.50 %, 1.75 % and 0.6 % respectively.

### 2.3 Experimental set-up

#### Dyadic set-up

Participants were comfortably seated in front of a square table with the lower arm resting on the table. The actor was seated at the left side of the table (seen from the participant’s point of view) so that the participant and actor were angled 90 degrees towards each other (Figure 1C). The grip-lift manipulandum (‘inverted T-shape’) was placed in between both individuals and positioned so that both individuals could grasp and lift the inverted T-shape comfortably. Participant and actor were asked to place their hands on a predetermined location on their side of the table to ensure consistent reaching throughout the entire experiment. Reaching distance was approximately 25 cm and required both individuals to use their entire right upper limb to reach for the inverted T-shape. When the actor would execute a trial, he/she would reach with their arm in front of the participant’s upper body and lift the manipulandum from this position. We opted for placing actor and participant side by side (and not opposite) for two reasons: First, Mojtahedi et al. (2017) demonstrated that, when executing a dyadic interaction task simultaneously, subjects produce a smaller lifting error in a side-by-side configuration compared to a face-to-face one. Second, from this position, observed actor lifts had the same frame of visual reference as the lifts executed by the participants. Arguably, this would enhance participant’s performance as it has been shown that corticospinal excitability during action observation is increased when observing actions from a first person point of view (Alaerts et al. 2009; Gallese et al. 2004). A transparent switchable screen (MagicGlass) was placed in front of the participant’s face which was transparent during trials and returned opaque during inter-trial intervals. The switchable screen ensured that participants could not see the experimenter switching the cuboids between compartments, thus making them ‘naïve’ to the actual center of mass. Last, trials lasted four seconds and their onset was indicated with a neutral sound cue and the switchable screen (and glasses; see below) turning transparent. Trial length ensured that participant and actor had enough time to reach, grasp, lift and return the object smoothly at a natural place. Inter-trial interval was approximately five seconds during which the screen returned opaque and the center of mass could be changed.

#### Experimental groups

As mentioned above, participants were assigned to either the ‘skilled observation group’ or the ‘error observation group’. Participants in the error observation group were paired with another participant and served as actors for each other (see: ‘Experimental procedure’). To ensure that the ‘participant actors’ were also naïve to the center of mass change, they were required to wear transparent switchable glasses (PLATO, Translucent technologies) which behaved identically as the switchable screen. Participants in the skilled observation group were paired with the experimenter who also served as the actor. For the skilled observation group, the ‘experimenter-actor’ was seated, as described above, on the left side of the participant (thus leaving the experimenter seat in Figure 1C vacant). One of the authors (G. Rens) served as the actor for all participants in the skilled observation group. In the error observation group, the experimenter was seated opposite to the participant (and left of the actor; Figure 1C) as he still needed to change the center of mass between trials. However, he did not participate in the experimental task of the error observation group. During the experiment, the experimenter was also responsible for changing the object’s weight distribution (i.e., switching the cuboids). During this cube switching action, the experimenter (thus the informed actor for the skilled observation group) would lift and rotate the manipulandum to switch the cuboids. Afterwards he would place the manipulandum in the same orientation, requiring him again to lift the manipulandum. This would provide him with experience on the actual weight distribution each time as the informed actor.

### 2.4 Experimental procedure

#### General procedure

All participants performed the experimental task in two separate sessions with at least 24 hours between sessions. Each dyad (of both groups) performed both experimental sessions maximally five days apart (i.e., within the same week). During the first session, participants gave written informed consent and were explained the experimental task and received the following instructions regarding object lifting: (1) lift the inverted T-shape to a height of approximately five cm at a smooth pace that is natural to you. (2) Only use your right thumb and index finger and only place them on the graspable surfaces (see: ‘Data acquisition’). (3) You are free to position your fingers on the graspable surfaces according to your own preferences and regardless of the actor’s positioning in the previous trial. (4) Keep the inverted T-shape’s base as horizontal as possible during lifting (i.e., ‘try to minimize object roll’). (5) The center of mass in your trials always matches the one in the actor’s preceding trial. In sum, participants were explained that they should try to estimate the center of mass during observed lifting and subsequently try to minimize object roll during their own lifts. Importantly, participants were explicitly explained they were free to select their own digit positioning. Arguably, with these instructions, participants should develop a digit force-position coordination strategy according to their own preferences.

After task instructions, participants were given three practice trials for the symmetrical weight distribution and six practice trials for each asymmetrical distribution (left or right). For the practice trials on the center of mass, participants were asked to always place the fingertips at the same height as it is not possible to minimize object roll with the fingertips positioned at different heights. In half of the practice trials for asymmetrical weight distribution, participants were asked to place their fingertips on the same height, i.e., ‘collinear’ positioning. In the other half, they were asked to place their fingertips at different heights, i.e., ‘noncollinear’ positioning (left center of mass: right thumb higher than right index finger; right center of mass: right thumb lower than right index). We emphasized these two different digit positioning on the asymmetrical weight distribution to ensure that participants would understand the full scope of possibilities for minimizing object roll. As we mentioned that participants were free position their fingertips according to their own preferences, these practice trials were intended to have participants understand there were multiple, equally appropriate, ways of positioning the fingertips and minimize object roll. Last, task instructions and practice trials were repeated at the start of the second session.

#### Experimental task

After task instructions, participants performed the object lifting task in turns with the actor. Actor and participant alternatingly performed a pseudo-random amount of three to six lifts on the same center of mass (for an example see Figure 2A but please note that the example shows maximally two repetitions per weight distribution). Accordingly, the length of a ‘sequence’ (i.e., sequential lifts on the same center of mass; Figure 2A) varied between an even amount of six and 12 lifts (six and 12 included; instead of the four shown in Figure 2A). After a sequence was completed, the experimenter changed the center of mass for the next sequence, i.e., a ‘transition’ from one weight distribution to the other (Figure 2A). Due to the even number of lifts per sequence, the person in the actor role always lifted the new center of mass first. To ensure that participants (and naïve actors) could not rely on sound cues (related to changing the center of mass) to locate the new center of mass, the experimenter always removed and replaced all three cubes after randomly rotating the inverted T-shape prior to each actor trial. These actions were never done before participant trials as they were explained that the center of mass in their trials would always match the one of the actor’s preceding trial.

**Figure 2.**
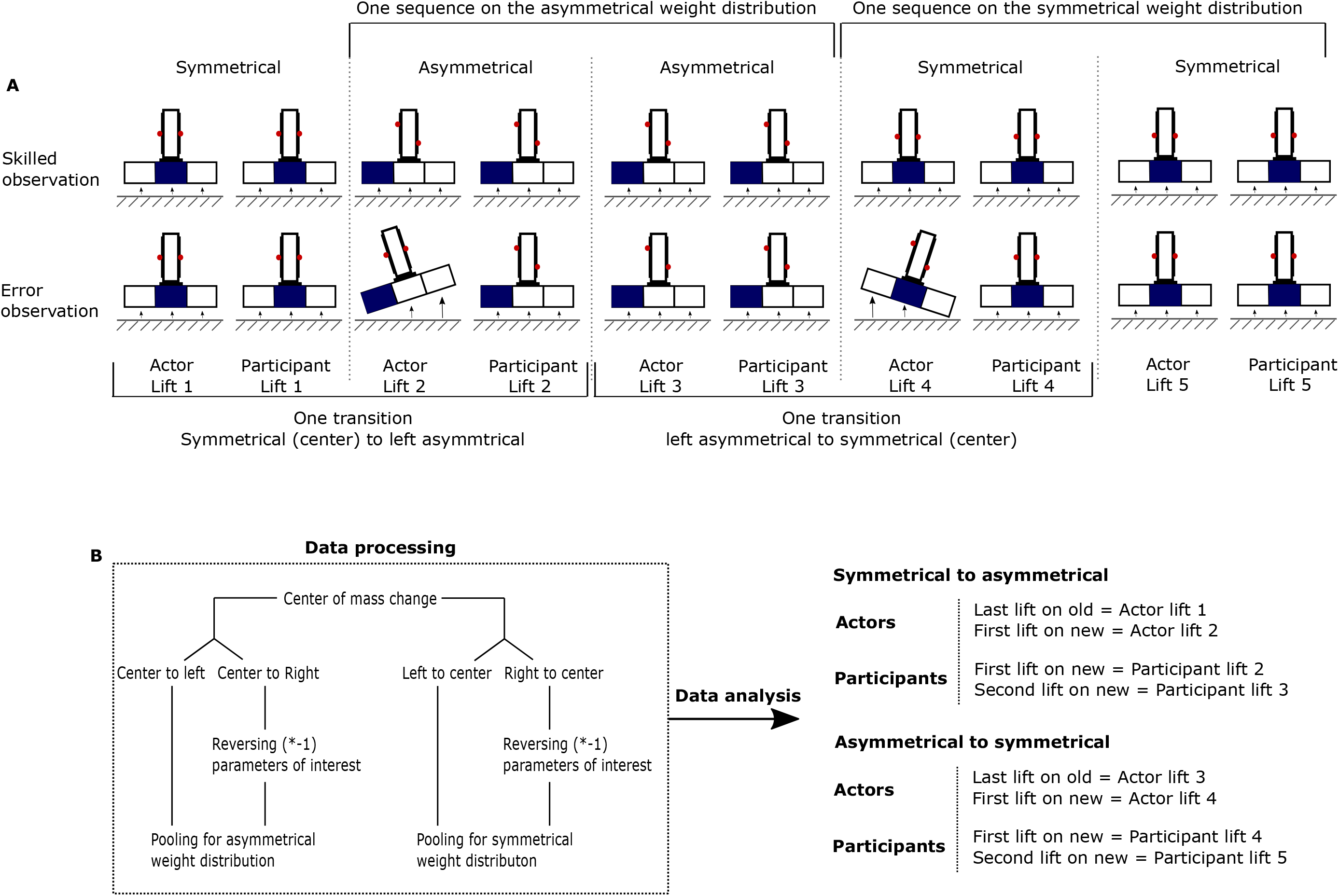
Lifting sequence and data analysis. **A**. Example of object lifting sequence of a subject and their respective actor (top row: skilled observation group with an informed actor; bottom row: error observation group with a naïve actor). The manipulandum’s base consisted of three compartments in which differently weighted cubes could be placed. The blue compartment indicates the location of the heavy cube whereas the white compartments indicate the location of the light cubes thus causing symmetrical (middle compartment is blue) and asymmetrical weight distributions (one compartment on the side is blue; left compartment in Figure 1A). Red dots indicate examples of index (left) and thumb (right) positioning when lifting the manipulandum. A sequence is defined as the actor and participant lifting the same weight distribution repeatedly. A transition is defined as the actor and participant having lifted the old weight distribution (e.g., symmetrical; actor lift 1 and participant lift 1) and then lifting the new weight distribution (e.g., asymmetrical; actor lift 2 and participant lift 2). **B**. Structure of how data was collected separately for the weight distributions, how it was subsequently pooled and analyzed for actors and participants separately.

#### Experimental task in the skilled observation group

During the alternating task, the informed actor and participant performed 20 transitions from the center of mass to each side. The experimenter lifted 10 center of mass sequences on each side with his fingertips placed collinearly and 10 with his fingertips placed noncollinearly. Importantly, for the informed actor, digit positioning was not marked in order to not bias participants in their own digit positioning. However, before onset of the study, the informed actor practiced the collinear and noncollinear lifts to ensure systematic lift performance. Moreover, for the noncollinear condition, the informed actor always attempted to position his fingertips as wide apart as possible, while still being able to lift the manipulandum comfortably. This was done to ensure maximal clarity of visual cues related to digit positioning.

We decided on 10 sequences per condition based on Reichelt et al. (2013) who used eight sequences. We included two more to take potential errors of the actor in account. We argued that experimentally manipulating the experimenter’s digit positioning would enable us to investigate whether participants rely upon observed digit positioning to perceive the object’s weight distribution. Importantly, as the experimenter was responsible for changing the center of mass, he should have always lifted the inverted T-shape skillfully. As such, participants had to rely on other lifting parameters (such as digit positioning) to perceive the object’s center of mass. After the sequence on the left or right side was completed, the experimenter changed the center of mass back to the center position to ‘wash out’ the internal representation for the asymmetrical weight distribution (for example see Figure 2A; Actor lift 3 and Actor lift 5). In addition, 10 ‘catch transitions’ in which the center of mass changed from side to side (and not side to center) were included to ensure that participants would not anticipate the typical change from side to center compartment. Transition orders were pseudo-randomized for each participant.

Importantly, the skilled observation group also performed the lifting task without actor to assess baseline sensorimotor memory effects (for example see: Johansson and Westling, 1984). This condition was included to investigate the magnitude of the lifting errors participants would make in the absence of lift observation. In this ‘without observation’ condition participants performed 10 transitions from the center to each side and five catch transitions. Similar to the alternating task, the experimenter changed the center of mass every third to six trial performed by the participant. The alternating (with actor) and without observation condition (without actor) were split over four and two experimental blocks, respectively. Participants performed two alternating and one without observation block in one session and the other blocks in the second session. Participants started or ended one session with the without observation block to counter-balance order effects across participants. Participants received a short break between blocks. Last, the lifting sequences were equally distributed over all blocks. That is, each of the four blocks of the alternating lifting task consisted of five center of mass transitions to each side. In addition, in each block and for each side, participants observed the actor using each digit positioning type either two or three times (due to 10 trials per condition not being divisible over four blocks).

#### Experimental task in the error observation group

In addition to general instructions, participants in the error observation group were also explained that both of them would perform the participant and actor roles. Each participant performed 10 center of mass changes from the center to each side position and five catch transitions in each role, i.e., once as actor and once as participant (Figure 2A bottom row). As such, participants performed 20 ‘experimental’ and 10 catch transitions for both roles combined. As actors were naïve to the center of mass change, they could not anticipate the center of mass change causing them to not generate the appropriate amount of compensatory torque and having the object roll towards its heavy side. Importantly, because actors in this group were naïve, we could not experimentally manipulate their digit positioning as we did in the skilled observation group. Because of this, the error observation group observed only half the amount of transitions compared to the skilled observation group [error observation group: 10 center to each side transitions; skilled observation group: 20 center to each side transitions (10 for collinear digit placement and 10 for noncollinear digit placement condition)]. Last, we did not include a without observation condition in the error observation group as the trials of participants in the actor role could be used as the ‘without observation condition’ to investigate baseline sensorimotor memory effects (for example see: Reichelt et al. 2013). To end, participants of the error observation group performed four experimental blocks of which two as actor and two as participant, spread over two sessions. Participants received short breaks between each block. Each participant started one session as actor and the other one as participant, switching seats within sessions when changing roles. Before participants performed as actor, they were given the same practice trials to get familiarized with their new seating. Due to this set-up, participants knew that the center of mass change would always happen first to the participant in the actor role. Accordingly, ‘actor participants’ were also explicitly asked to not guess or try to predict the center of mass change. During breaks and until the end of the second session, participants were not allowed to discuss the experiment with each other.

### 2.5 Data analysis

Data collected with the F/T sensors and accelerometer were sampled in 3 dimensions at 1000 Hz and smoothed using a fifth-order Butterworth low-pass filter (cut-off frequency: 15 Hz). On each force sensor, grip force (GF) was defined as the exerted force perpendicular to the normal force (Y-direction on Figure 1) and load force (LF) was defined as the exerted force parallel to the normal force (X-direction on Figure 1). Digit center of pressure was defined as the vertical coordinate (X-direction on Figure 1) of the center of pressure of the finger on the graspable surface attached to each force sensor and was calculated from the force and torque components measured by the respective F/T sensor relative to its frame of reference. For each sensor, the center of pressure was computed with formula 1. 

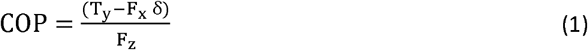

In formula 1, COP = center of pressure; T_y_ = Torque in the Y-direction, F_x_ = Force in the X-direction, F_z_ = Force in the Z-direction, δ = cover plate thickness (1.55 mm). Using the digit center of pressure, we could also compute the compensatory torque. Compensatory torque was defined as the net torque generated by an individual to offset the external torque (i.e., to minimize object roll) caused by the object’s asymmetrical weight distribution. Compensatory torque was computed with formula 2 (we refer the reader to the supplementary materials of Fu et al., 2010 for the explanation of the formula). 

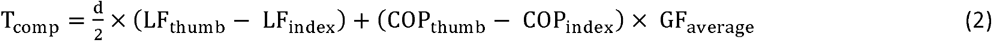

In formula 2, T_comp_ = Compensatory torque, d = horizontal distance between the digits (48 mm; Figure 1; Y-direction), LF_thumb/index_ = Load force generate by the thumb and index finger respectively, COP_thumb/index_ = center of pressure of the thumb and index finger respectively, GF_average_ = averaged amount of GF exerted by the thumb and index finger. Object roll was measured using the accelerometer and was defined as the amount of tilt over the X-axis, thus in the Y-direction (same directionality as T_comp_ depicted on Figure 1B). Object roll was computed with formula 3. 

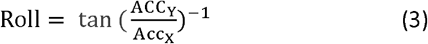

In formula 3, Roll is the object roll as described above, tan^-1^ is the inverse tangent, or arctangent, ACC_Y_ and ACC_X_ are the acceleration in the X- and Y-axes respectively (Figure 1B).

To investigate the effects of lift observation on the participants’ performance, we used the following variables: Object roll, digit positioning difference, defined as the difference between the COP of the thumb and the index finger (positive values indicate a thumb placement higher than that of the index finger) and compensatory torque. We included difference in digit positioning to investigate whether the error and skilled observation groups used a different digit positioning strategy after lift observation. In addition, we emphasized object roll as our key indicator of performance as (a) it reflects the instructed task requirement (i.e., ‘minimize object roll’) and (b) because object tilt is visible during lift information and could potentially convey critical information about the weight distribution to the participants. Compensatory torque was included to investigate the participants’ lift planning and because Fu et al. (2010) showed high correlations between object roll and compensatory torque. For the actors, we also included total grip force and load force difference (analogue to digit positioning difference; LF thumb minus LF index finger) at lift onset to explore potential differences in observed lift performance. In line with Fu et al. (2010), digit positioning was extracted at early object contact and the other parameters (compensatory torque, load force difference and total grip force) were extracted at lift onset. Early object contact was defined as total GF > 1 N to ensure that proper contact was established with both fingertips. Lift onset was defined as the first peak in the vertical acceleration (X-direction on Figure 1) between object contact and 200 ms after object lift off (defined as total LF = object weight). Last, in line with Fu et al. (2010), peak object roll was defined as the maximum object roll to either side (and thus could be positive or negative) between object contact and 200 ms after object lift off.

For each participant and each condition separately, we calculated the means of the lifting parameters that were used in the analyses described below. Accordingly, for a given lifting parameter and condition, we had 12 data points for the error observation group and 11 data points for the skilled observation group. Because we did not expect relevant differences for lift performance between the left and right asymmetrical weight distribution (i.e., similar values for compensatory torque on either weight distribution). In addition, we also did not expect relevant differences for the symmetrical weight distribution depending on whether the previously lifted weight distribution was left or right asymmetrical. Moreover, we were not interested in whether lift performance differed between the left and right sides, but rather whether lift observation can improve predictive lift planning when lifting an unexpected asymmetrical or symmetrical weight distribution. In addition, potential statistical differences between sides could be caused by the hidden multiplicity of multiple testing (Cramer et al., 2016). Because of these reasons, we decided to pool our data for ‘side’. As shown in Figure 2B, we pooled lift performance for the asymmetrical weight distributions (center of mass change to left or right) and we pooled for the symmetrical weight distribution (center of mass change from left or right). To ensure that pooling would not eradicate all effects (e.g., compensatory torque generated clockwise or counterclockwise is positive and negative respectively; Figure 1), we reversed the sign for object roll, compensatory torque, digit positioning difference and load force difference for when the center of mass changed to or from the right side (Figure 2B). For the two latter parameters, positive values for the respective parameters indicate that (1) the finger on the heavy side is positioned higher than the finger on the light side and (2) the finger on the heavy side generates more load force than the finger on the light side. Importantly, uniquely for the symmetrical weight distribution, positive values indicate that the fingertip, previously on the heavy side, is positioned higher/generates more load force than the fingertip that was previously on the light side.

### 2.6 Statistical analysis

When the center of mass is on either side (i.e., ‘asymmetrical weight distribution’), participants need to generate the appropriate amount of compensatory torque to offset the external torque and minimize object roll. In contrast, when the center of mass is in the middle (i.e., ‘symmetrical weight distribution’), participants need to minimize object roll by not exerting any compensatory torque. Considering that these ‘task goals’ are different (asymmetrical: generate compensatory torque; symmetrical: do not generate compensatory torque), we analyzed lift performance separately for the asymmetrical and symmetrical weight distributions. All statistical analyses were performed in SPSS statistics version 25 (IBM, USA) and are described below.

#### Lifting performance of the actors

To investigate potential differences in the actors’ lifting performance, we performed the following analysis on each of the five included lifting parameters separately. For the actors, we included their third lift on the ‘old’ weight distribution and their first lift on the ‘new’ unexpected weight distribution. For example, when the center of mass would change from symmetrical to asymmetrical, we would take Actor lift 1 (‘old’ symmetrical weight distribution) and Actor lift 2 (‘new’ asymmetrical weight distribution) as depicted in Figure 2A. Accordingly, when the center of mass would change from asymmetrical to symmetrical, we would take Actor lift 3 and Actor lift 4 as depicted in Figure 2A. In line with Rens and Davare (2019), we did not use the last lift, but the third one. As the actors (and participants) would lift a given weight distribution three to six times, we decided to use the last lift of the consistent repetition (third lift) rather than the actual last lift. This approach allowed us to investigate whether actors would appropriately update their predictive object lifting command for the novel weight distribution by lifting the new weight distribution differently than the old one. We did not include the actors’ second lift on the new weight distribution as the observing participants might have only relied on lift observation to plan their first lift and afterwards relied entirely on haptic feedback from their previous lift to plan upcoming lifts. For instance, when the weight distribution would change to asymmetrical, we included Actor lift 1 and Actor lift 2, but not Actor lift 3 in Figure 2A.

Considering that we had an incomplete factorial design based on our experimental groups, we decided to use linear mixed models (LMMs). We included the factor GROUP (skilled or error observation groups), DISTRIBUTION (old weight distribution and new weight distribution) and OBSERVATION (skilled noncollinear, skilled collinear, error observation). We included GROUP and DISTRIBUTION as main effects as well as their interaction effect GROUP X DISTRIBUTION. Due to each group observing only one lifting performance type (skilled lifts or lifting errors), OBSERVATION was added as a factor nested within GROUP (i.e., OBSERVATION_GROUP_). Last, we also included the interaction effect DISTRIBUTION X OBSERVATION_GROUP_

#### Lifting performance of the participants

To investigate the participants’ lifting performance, we included their first and second lift on the new, unexpected weight distribution. For instance, when the center of mass changed from symmetrical to asymmetrical, we would take Participant lift 2 and Participant lift 3 of the example depicted in Figure 2A. Accordingly, when the center of mass changed to symmetrical, we would take Participant lift 4 and Participant lift 5 of the same example depicted in Figure 2A.

In line with Rens and Davare (2019) and Reichelt et al. (2013), the potential effects of lift observation on predictive lift planning can be investigated by comparing lift performance after lift observation with lift performance after having haptic feedback about the actual object properties. Accordingly, here we were not interested in whether participants plan their lift differently for the new weight and old weight distribution, but rather whether they plan their lift for a new weight distribution similarly after lift observation or haptic feedback. Similar to our analyses for the actors’ lift performance, we included the same factors GROUP and OBSERVATION. Importantly, there are two major differences with the actors’ analyses. First, the factor DISTRIBUTION has been termed ‘REPETITION’ here (first or second lift on the new weight distribution), as both included lifts were performed on the same weight distribution. Second, the factor OBSERVATION consists of four levels here, i.e., skilled noncollinear, skilled collinear, error observation and also ‘without observation’). Accordingly, we included GROUP and REPETITION as main effects as well as their interaction effect GROUP X REPETITION. Due to each group observing only one lifting performance type (skilled lifts or lifting errors), OBSERVATION was added as a factor nested within GROUP (i.e., OBSERVATION_GROUP_). Last, we also included the interaction effect REPETITION X OBSERVATION_GROUP_. Last, we did not include the third lift on the novel weight distribution as Fu et al. (2010) showed that predictive lift planning on an object with unexpected weight distribution improves from the first to second lift but not anymore from the second to third one which was supported by our preliminary analyses including this lift.

Importantly, because actors and participants performed two separate sessions (both groups) and switched roles (only error observation group), we investigated day and order effects. As these preliminary analyses did not reveal any relevant significant differences, we decided to pool our data across sessions and experimental blocks. Finally, for all LMMs, we decided to include the mixed model covariance structures as first-order autoregressive based on the assumption that the correlation in residuals between factor levels was identical across levels. For the actors’ LMMs we included the actors as subjects in the model (same actor for the skilled observation group and 12 unique actors for the error observation group). For the participants’ LMM we included the participants as subjects in the model (11 participants in the skilled observation group and 12 participants in the error observation group). In each LMM, we also included the intercept. We used type III sum of squares and Maximum Likelihood (ML) for mixed model estimation and Bonferroni for pairwise comparisons. All data is presented as the mean ± standard error of the mean (SEM).

Last, Fu et al. (2010) reported high correlations (r > 0.89) between compensatory torque at lift onset and peak object roll in their study. To validate findings in our study, we calculated Pearson correlations between peak object roll and compensatory torque at lift onset for our participants as well. Correlations were calculated separately for the weight distribution changes to symmetrical and to asymmetrical. All participants were taken together (23 participants across both groups) and we included both the first and second lifts after the weight distribution change for all conditions (for example, for the change to asymmetrical these trials would be Participant lift 3 and Participant lift 4 in Figure 2A). Specifically, for the error observation group we included the error and without observation conditions and for the skilled observation group we included the collinear, noncollinear and without observation conditions. To end, we did not enter separate trials, but the individual participant means for each condition.

## 3. Results

In the present study, we aimed to investigate whether individuals are able to perceive an object’s weight distribution during observation of object lifting and subsequently use this information to update their own predictive lift planning. Participants performed an object lifting task in turns with an actor and were asked to lift the object as skillfully as possible (i.e., minimize object roll by generating the appropriate amount of compensatory torque). Conversely, lifting was performed erroneously when an individual exerted an incorrect amount of compensatory torque causing the object to roll, i.e., ‘tilt’, towards its heavy side during lifting. Participants were separated over two groups and paired with specific actors. The skilled observation group was paired with the experimenter (‘informed actor’) who could anticipate the center of mass change and would lift the object skillfully. Participants in the error observation group were paired with other participants from this group and served as actors for each other. The actors in the error observation group could not anticipate the center of mass change and were thus ‘naïve’ (‘naïve actors’). Finally, participants could potentially rely on observed lifting to estimate the object’s weight distribution and subsequently plan their own lifts correctly.

In Figure 3, traces of the different lifting parameters are shown. Each color represents the parameter traces of one exemplary lift. Green and blue traces represent single lift examples when skillfully lifting an asymmetrical weight distribution with noncollinear and collinear digit positioning respectively. Red traces represent the single lift example when erroneously lifting the asymmetrical weight distribution with collinear digit positioning.

**Figure 3.**
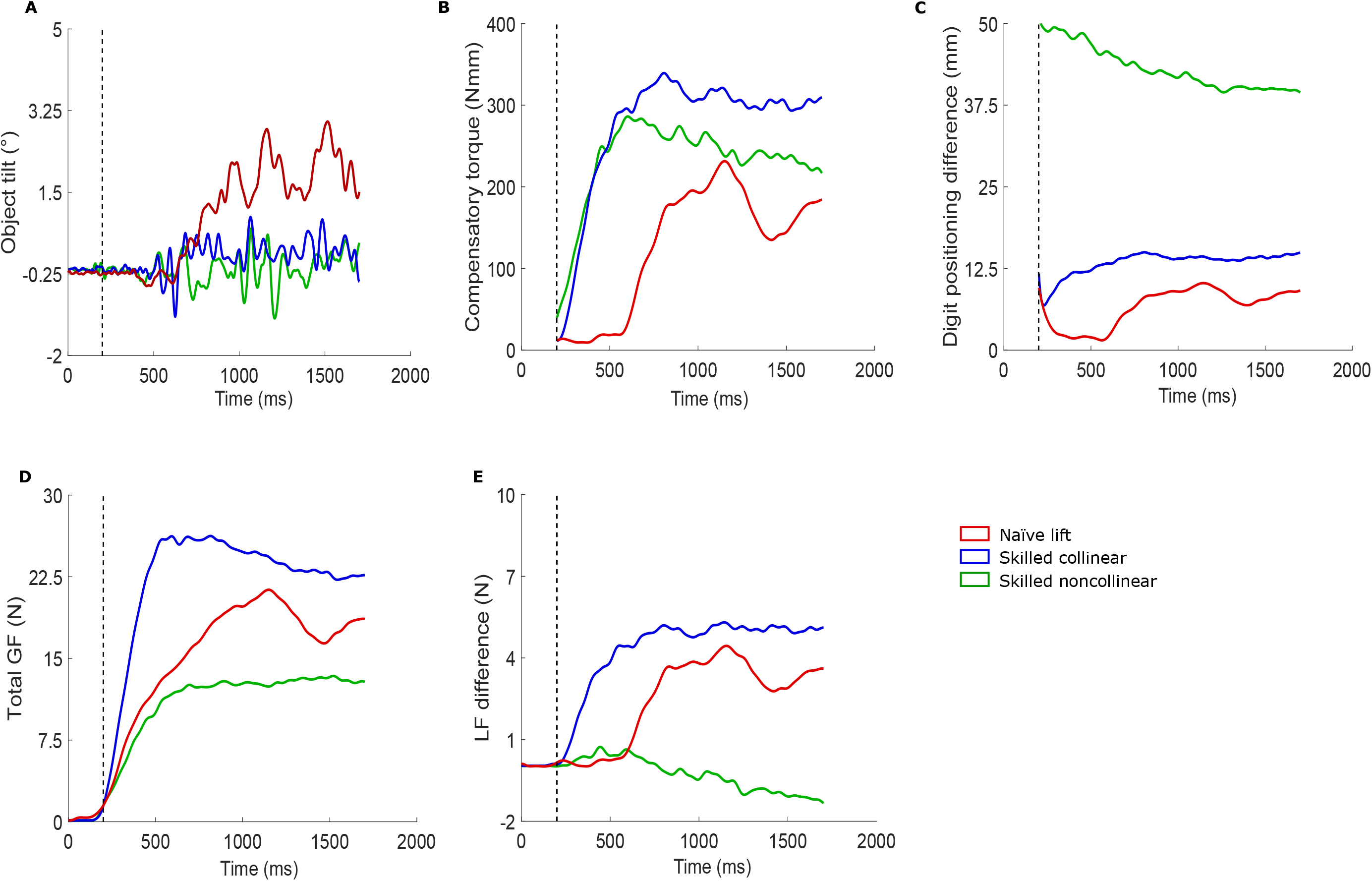
Representative traces when lifting the asymmetrical weight distribution for each experimental condition. Typical traces showing the evolution of different parameter profiles over time for a skilled lift with fingertips positioned on the same height (‘skilled collinear’), a skilled lift with fingertips positioned on different heights (‘skilled noncollinear’) and a naïve lift (‘naïve lift’) in which the individual incorrectly anticipated the weight distribution to be symmetrical. **A**. Amount of object tilt (°) caused by the participant (not) generating appropriate compensatory torque. **B**. Compensatory torque (Nmm) generated by the individual to offset the external torque, induced by the asymmetrical weight distribution. **C**. Digit positioning difference (mm): vertical distance between the centers of pressure of the fingertips (position fingertip on heavy side - position fingertip on light side). **D**. Total amount of grip force (N). **E**. Difference in load force exertion (N) between the fingertips (load force fingertip on heavy side – load force fingertip on light side). The dashed black line represents early object contact. As compensatory torque and digit positioning difference are calculated based on early contact and are highly contaminated by noise before actual contact, we removed their values before early object contact.

When an individual correctly plans to lift an asymmetrical weight distribution, they will induce limited object tilt (Figure 3A blue and green traces; i.e., ‘skilled lifts’). Importantly, object tilt is minimized as the participant predictively generates appropriate compensatory torque (Figure 3B blue and green traces) to offset external torque induced by the object’s weight distribution. In contrast, when an individual expects a symmetrical weight distribution, they will not predictively generate compensatory torque for the actual asymmetrical weight distribution (Figure 3B red trace, i.e., ‘‘lifting error’) which causes object tilt (Figure 3A). As such, an individual’s expectations of the object’s weight distribution can be probed by quantifying the amount of object tilt and compensatory torque they generated.

Importantly, generating compensatory torque relies on generating a valid digit positioning – (load) force coordination pattern (for a detailed explanation see Fu et al. 2010). The blue traces on Figure 3C and 3E resembles an individual skillfully lifting an asymmetrical weight distribution when placing the fingertips on the same height. As the vertical height difference between the fingertips is small (Figure 3C blue trace), the fingertip on the heavy side has to generate more load force than the finger on the light side (Figure 3E blue trace). Conversely, the green traces on figure 3C and 3E resembles an individual skillfully lifting an asymmetrical weight distribution with noncollinear positioning (i.e., fingertip on heavy side positioned higher than fingertip on light side). Due to the vertical height difference between the fingertips (Figure 3C green trace), a ‘lever’ is created which results in the fingertip on the heavy side having to generate less load force than in the collinear condition. As a result, the load force difference in the noncollinear condition (Figure 3E green trace) becomes much smaller than in the collinear condition (Figure 3E blue trace). Last, when an individual incorrectly expects a symmetrical weight distribution, they will position their fingertips on the same height (Figure 3C red trace). As a result, the fingertip on the heavy side should generate more force than the one on the light side. However, as they incorrectly planned their lift, this load force difference between the fingertips will be generated slower compared to when they correctly anticipated the asymmetrical weight distribution (i.e., the difference in load force difference between the blue and red trace in Figure 3E which is analogue to the difference for compensatory torque in Figure 3B). Last, as can be seen in Figure 3D, the fingertips generate more grip force when positioned collinearly compared to noncollinear positioning. Analogue to compensatory torque and load force difference, grip force scaling is slower when performing a lifting error with collinear positioning (Figure 3D red trace) than when lifting skillfully with the same digit positioning (Figure 3D blue trace).

### 3.1 Center of mass change from symmetrical to asymmetrical

#### 3.1.1 Actor’s lifting performance

Based on the difference in naivety between the informed and naive actors, we expected that the informed actor would lift the asymmetrical weight distributions significantly better than the naive actors. As mentioned before, we included their last (i.e., third) lift on the ‘old’ symmetrical weight distribution and their first lift on the ‘new’ asymmetrical one. Arguably, if the actors’ lifting parameters when lifting the asymmetrical weight distribution were similar to those when lifting the symmetrical one, this would indicate that the actors would predictively plan their lift for the symmetrical weight distribution, thus erroneously for the new asymmetrical one. Although we included only the first lift on the new weight distribution, please note that the naive actors’ performance is the same as the error observation group’s performance in the ‘without observation condition’. Accordingly, the naïve actors’ repeated performance (first and second lifts) on the asymmetrical weight distribution could be found there. To end, for the actors’ performance we included (1) object tilt, as primary proxy of lift performance (Fu et al., 2010), (2) compensatory torque, which is used to offset the object’s external torque and minimize object tilt, (3) digit positioning, for investigating the informed actor’s compliance with task instructions, and (4) total grip force and (5) load force difference as these force parameters have been considered indicative of object weight (Alaerts et al., 2010; Hamilton et al., 2007) and could potentially convey similar information about an object’s weight distribution. Last, the actor’s lifting performance can be found in Figure 4.

**Figure 4.**
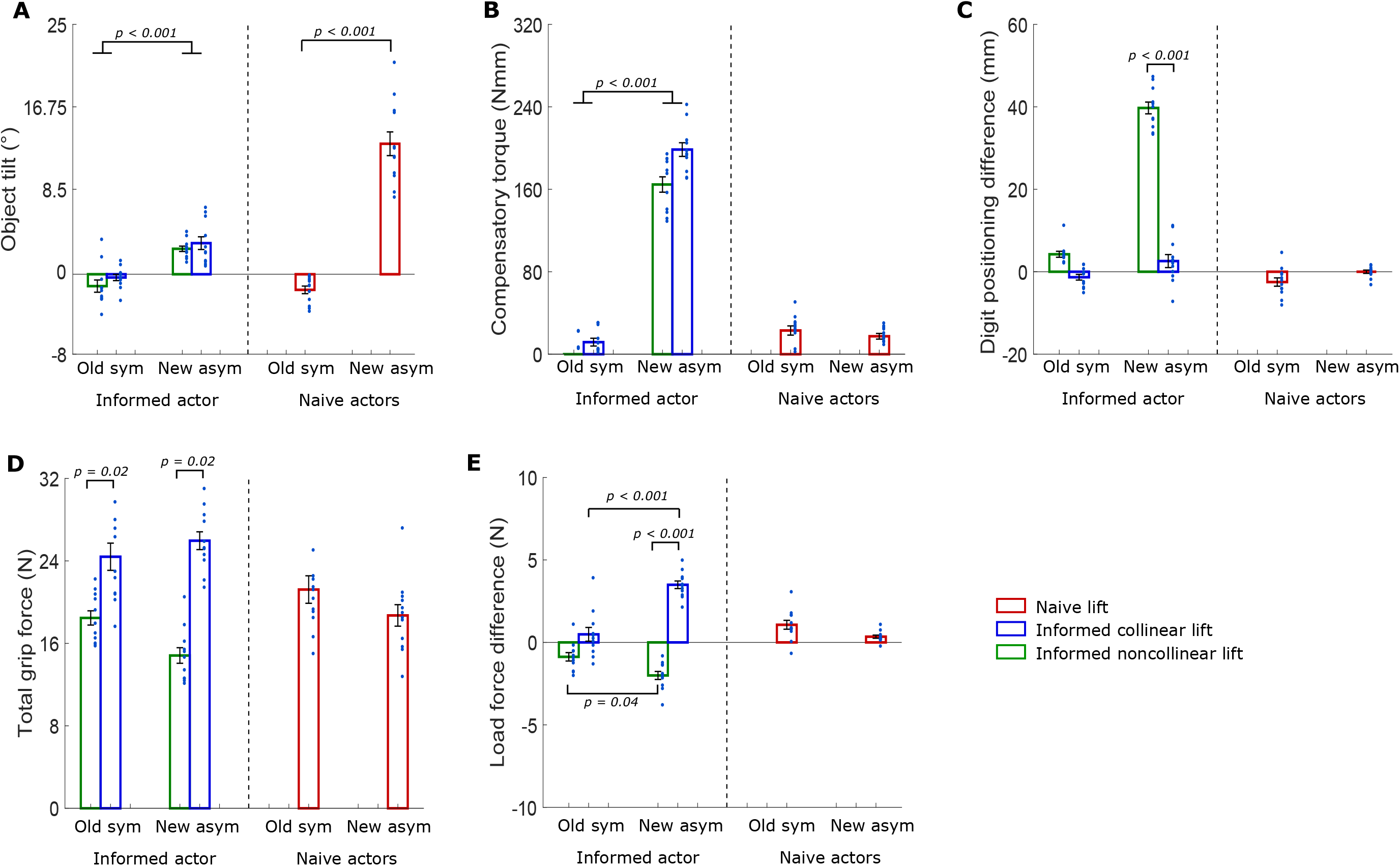
Lift performance of the actors. Lifting performance of the informed (n = 11, 8 females, for all conditions) and naïve actors (n = 12, 9 females, for all conditions) when the weight distribution changed from old symmetrical (‘Old sym’) to new asymmetrical (‘New asym’). Accordingly, the first bars (above ‘Old sym’) represent the actors’ third lift on the old symmetrical weight distribution and the second bars (above ‘ New asym’) represent the actors’ first lift on the new symmetrical weight distribution. **A**. Object tilt (°) **B**. Compensatory torque (Nmm). **C**. Digit positioning difference (mm). **D**. Total amount of grip force (N). **E**. Difference in load force exertion (N). **Red**: Lifting performance of the naïve actors who could not anticipate the weight distribution change (‘Naïve lift’). **Blue**: Lifting performance of the informed actor, who could anticipate the weight distribution change, when positioning both fingertips on the same height when lifting both weight distributions (‘Informed collinear lift’). **Green**: Lifting performance of the informed actor when positioning both fingertips on the same height on the symmetrical weight distribution but on different heights when lifting the asymmetrical one (‘Informed noncollinear lift’). All data is presented as the mean ± SEM. Each circle (scatter) represents the lift performance of one actor in a given condition. Only significant within-actor group differences are shown on the figure. Linear mixed models with Bonferroni post-hoc comparisons were used for the analyses.

#### Object tilt

Because, we are primarily interested in lifting performance differences between actor types on both weight distributions, we will discuss these findings in light of the significant effect DISTRIBUTION X GROUP (*F*_*(1,136)*_ *= 20.49, p < 0.001)*. As can be seen in Figure 4A, we did not find any evidence that the informed actor and naïve actors generated different amounts of object roll when lifting the old symmetrical weight distribution (informed old symmetrical = −0.76 ± 0.35 °; naive old symmetrical = −1.56 ± 0.38 °; *p = 1.00*). When the weight distribution changed to asymmetrical, the naïve actors induced more object roll (new asymmetrical = 13.05 ± 1.20°; *p < 0.001*), compared to their last lift on the old weight distribution (Figure 4A). Importantly, this indicates that the naïve actors were not able to anticipate the unexpected weight distribution change. However, it is important to note that the informed actor also induced more object roll in his first lift on the new asymmetrical weight distribution (mean = 2.83 ± 0.34 °; *p < 0.001*). However, on the new asymmetrical weight distribution, the informed actor induced significantly less object roll than the naïve actors *(p < 0.001)*. In sum, these findings substantiate the difference in naivety between actor types. That is, the informed actor was better at minimizing object roll after an (unexpected) change in weight distribution. Last, the effect DISTRIBUTION X OBSERVATION_GROUP_ *(F*_*(1,136)*_ *= 0.35, p = 0.56)* was not significant. Accordingly, these findings provide no evidence that the informed actor minimized object roll differently with collinear or noncollinear digit positioning.

#### Compensatory torque at lift onset

As can be seen in Figure 4B, the informed actor knew that the weight distribution would be asymmetrical, enabling him to increase the compensatory torque he generated (old symmetrical: mean = 2.90 ± 2.96 Nmm; new asymmetrical: mean = 181.59 ± 4.95 Nmm; *p < 0.001*) while the naïve actors, blind to this change, could not do so (old symmetrical: mean = 22.98 ± .47 Nmm; new asymmetrical: mean = 17.39 ± 2.77 Nmm; *p = 1.00*) (DISTRIBUTION X GROUP: *F*_*(1,136)*_ *= 43.26, p < 0.001*). As a result, the informed actor generated more compensatory torque for the asymmetrical weight distribution than the naïve ones (*p < 0.001*) substantiating their difference in naivety based on our experimental set-up. Last, we did not find any evidence that the informed actor generated different compensatory torque on the asymmetrical weight distribution when using collinear (mean = 198.54 ± 6.62 Nmm) or noncollinear positioning (mean = 164.58 ± 7.44 Nmm; *p = 0.59*) (DISTRIBUTION X OBSERVATION_GROUP_: *F*_*(1,136)*_ *= 0.20, p = 0.65*).

#### Digit positioning difference at early contact

We did not find any significant differences with respect to the digit positioning of the informed actor (mean = 1.45 ± 0.58 mm) and naïve ones (mean = - 2.50 ± 1.01 mm) when lifting the old symmetrical weight distribution *(p = 0.23) (DISTRIBUTION X GROUP:* F_(1,136)_ = 27.63, p < 0.001). Again, as the naïve actors could not anticipate the new asymmetrical weight distribution, our analyses failed to reveal significant differences between their digit positioning on the old symmetrical and new asymmetrical weight distribution (mean = 0.00 ± 0.40 mm; *p = 1.00*) (Figure 4C). With respect to the informed actor, our findings indicate that he complied with task instructions. In the collinear condition, he placed his fingertips not significantly different when using collinear positioning on the old symmetrical (mean = 1.45 ± 0.58 mm) and new asymmetrical weight distributions (mean = 2.59 ± 1.58 mm; *p = 1.00*) (DISTRIBUTION X POSITION_GROUP_: *F*_*(1,136)*_ *= 75.21, p < 0.001*). Conversely, when instructed to use noncollinear positioning on the new asymmetrical weight distribution, the informed actor placed his fingertips significantly further apart (mean = 39.73 ± 1.42 mm) compared to himself in all other conditions (*all p < 0.001)*.

#### Total grip force at lift onset

The grip forces the naïve actors generated when lifting the old symmetrical (mean = 21.22 ± 1.33 N) and new asymmetrical weight distributions (mean = 18.71 ± 1.04 N) did not differ significantly (*p = 1.00)* (Figure 4D). Similarly, when the informed actor was instructed to use collinear positioning (Figure 4D blue bars), he scaled his grip forces not significantly different when lifting the old symmetrical (mean = 24.41 ± 1.32 N) and new asymmetrical weight distributions (mean = 25.96 ± 0.86 N) *(p = 1.00)* (DISTRIBUTION X POSITION_GROUP_: *F*_*(1,136)*_ *= 4.42, p = 0.37)*. When the informed actor was instructed to change his digit positioning from collinear on the old asymmetrical weight distribution to noncollinear on the new asymmetrical one (i.e., the noncollinear condition; Figure 3C green bars), he also scaled his grip forces not significantly different for the old symmetrical (mean = 18.47 ± 0.69 N) and new asymmetrical weight distributions (mean = 14.82 ± 0.76 N; *p = 1.00)*. To end, the informed actor scaled his grip forces on average lower in the noncollinear (green bars) than in the collinear condition (*p = 0.02*) (POSITION_GROUP_: *F*_*(1,136)*_ *= 9.06, p = 0.003)*.

#### Load force difference at lift onset

As shown by the blue bars in Figure 4E, when the informed actor lifted the new asymmetrical weight distribution he scaled his load forces higher with the fingertip on the heavy side (mean = 3.49 ± 0.23 N) compared to when he lifted the old symmetrical weight distribution with the same digit positioning (mean = 0.49 ± 0.42 N; p < 0.001) (DISTRIBUTION X POSITION_GROUP_: *F*_*(1,136)*_ *= 26.36, p < 0.001)*. When the informed actor changed his digit positioning for the weight distributions, his load force difference was lower when lifting the new asymmetrical weight distribution noncollinearly (mean = −2.00 ± 0.24 N) compared to lifting the old symmetrical one collinearly (mean = −0.89 ± 0.25 N; *p = 0.04*) (Figure 3E green bars). Last, in line with our findings for the other lifting parameters, the naïve actors’ load force difference did not differ significantly when lifting the old symmetrical (mean = 1.07 ± 0.27 N) and new asymmetrical weight distributions (mean = 0.35 ± 0.09 N; p = 1.00) collinearly.

In sum, these lifting parameters substantiate our experimental set-up. That is, when the naïve actors repeatedly lifted the old symmetrical weight distribution (their third lift on the symmetrical weight distribution), their lifting parameters did not differ significantly from those in their first lift on the new asymmetrical weight distribution. Accordingly, when the naïve actors lifted the unexpected asymmetrical weight distribution for the first time, they were not able to minimize object tilt thus causing the object to roll towards its heavy side (Figure 4A). In contrast, as the informed actor could predict the weight distribution change, he anticipatorily scaled his lifting parameters significantly different when lifting the symmetrical and asymmetrical weight distributions (Figure 4B-4E). As a result, the informed actor induced significantly less object tilt when lifting the new asymmetrical weight distribution, compared to the naïve actors. However, the skilled actor still induced significantly more object roll compared to himself when lifting the old symmetrical weight distribution (Figure 4A).

#### 3.1.2 Participants’ lifting performance

To investigate potential improvements in predictive lift planning, we compared the participants’ lift performance in their first lift on the new weight distribution with their second ones. Logically, in their second lift participants could rely on haptic feedback. Thus, this comparison enabled us to investigate how lift planning based on observation compares to lift planning based on haptic feedback. The participants’ lifting performance on the asymmetrical weight distribution can be found in the top row of Figure 5.

**Figure 5.**
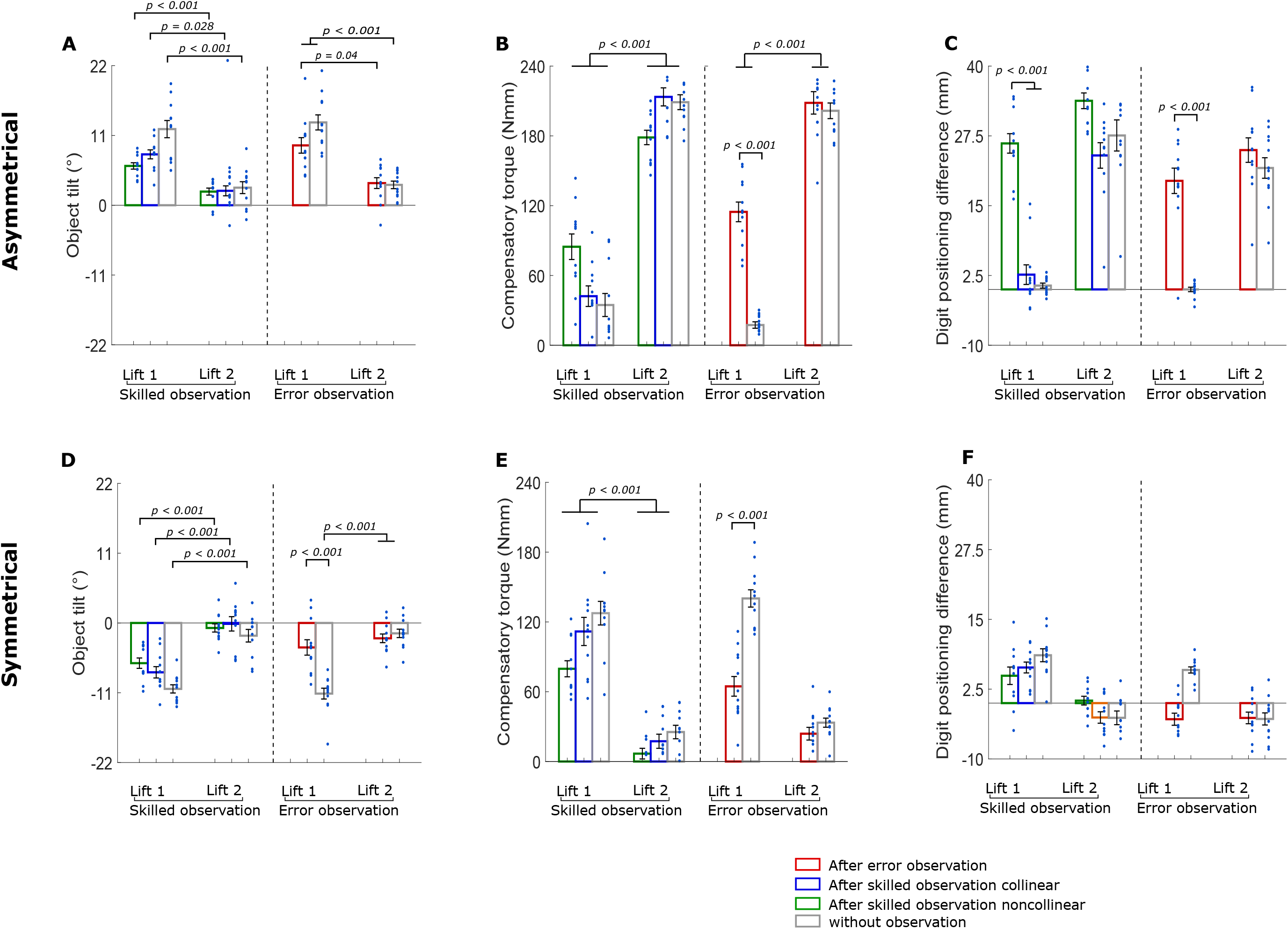
Lift performance of the participant. Lifting performance of the skilled (‘skilled group’; n = 11, 8 females, for all conditions) and error observation group (‘error group’; n = 12, 9 females, for all conditions) when the weight distribution changed from symmetrical to asymmetrical (**top row**) or from asymmetrical to symmetrical (**bottom row**). For each group first and second lifts on the novel weight distribution are shown. **A and D**. Object tilt (°) **B and E**. Compensatory torque (Nmm). **C and F**. Digit positioning difference (mm). **Red**: Lifting performance of the error group after observing a lifting error by the naive actors who could not anticipate the weight distribution change (‘error observation’). **Blue**: Lifting performance of the skilled group when observing the informed actor lifting both weight distributions with collinear digit positioning (‘skilled observation collinear’). **Green**: Lifting performance of the skilled group when observing the informed actor lifting the symmetrical and asymmetrical weight distributions with collinear and noncollinear digit positioning respectively (‘skilled observation noncollinear’). **Grey**: Lifting performance of each group when lifting without prior observation (‘without observation’). All data is presented as the mean ± SEM. Each circle (scatter) represents the lift performance of one participant in each condition. Only significant within-group differences are shown on the figure. Linear mixed models with Bonferroni post-hoc comparisons were used for the analyses.

##### Lift observation does not improve minimizing object tilt

Results regarding tilt can be found in Figure 5A. We will discuss these findings in light of the significant effect REPETITON X OBSERVATION_GROUP_ (*F*_*(3,228)*_ *= 3.73, p = 0.012)*. First, our findings provide no evidence that observing skilled lifts improve predictive lift planning. That is, the skilled observation group did not minimize object roll significantly better after observing a skilled lift (after noncolinear observation = 6.19 ± 0.50 °, Figure 5A first green bar; after collinear observation = 8.02 ± 0.71 °, Figure 5A first blue bar) than without observation (mean = 12.00 ± 1.35 °; Figure 5A first grey bar; *both p > 0.11*). In addition, in their second lifts (after noncollinear observation; Figure 5A second green bar; mean = 2.15 ± 0.55 °; after collinear observation; Figure 5A second blue; mean = 2.27 ± 0.78 °; without observation; Figure 5A second grey bar; mean = 2.77 ± 0.94 °), they minimized object roll significantly better than in their respective first lifts *(all p < 0.03)*. In sum, these findings indicate that observing skilled lifts did not improve the skilled observation group’s ability to minimize object roll without haptic feedback. Accordingly, the skilled observation group was only able to minimize object roll when having haptic feedback about the weight distribution (i.e., in their second lifts).

The error observation group, similar to the skilled observation group, performed significantly worse in their first lift after lift observation (Figure 5A first red bar; mean = 9.43 ± 1.22 °) than in their second one (Figure 5A second red bar; mean = 3.49 ± 0.85 °; *p = 0.04*). Moreover, our findings provide no evidence that the error observation group minimized object roll better in their first lift after lift observation than without it (Figure 5A first grey bar; mean = 13.05 ± 1.19 °; *p = 1.00*). When the error observation group could not anticipate the weight change they generated significantly more object roll (Figure 5A; first grey bar) compared to when they could rely on haptic feedback (Figure 5A; second grey bar; mean = 3.21 ± 0.59 °). Taken together, these findings indicate that observing lifting errors did not improve predictive object lifting and that the error observation group was only able to minimize object roll after having access to haptic feedback. To end, our analysis did not reveal any significant differences between the first lifts of the skilled and the error observation groups (*both p = 1.00)* as well as between the first lifts, after each skilled observation type, within the skilled observation group (*p = 1.00*). Accordingly, our findings provide no evidence that our lift observation types mediated predictive minimizing of object roll differently.

##### Error observation has a larger impact on predictive planning of compensatory torque at lift onset than skilled observation

As shown in Figure 5B, both groups exerted higher compensatory torques after having their internal object representation for the asymmetrical weight distribution updated through haptic feedback (second lifts pooled across conditions; mean = 202.33 ± 3.37 Nmm) compared to when they had no haptic feedback (first lifts pooled across conditions; mean = 59.77 ± 4.19 Nmm; p < 0.001) (REPETITION: *F*_*(1,228)*_ *= 636.67, p < 0.001)*. This effect of repetition is potentially driven by pooling of the ‘without observation’ and observation conditions. To investigate this possibility, we explored the significant effect of REPETITION X OBSERVATION_GROUP_ *(F*_*(3,228)*_ *= 16.78, p < 0.001)*. Interestingly, when the error observation group lifted the asymmetrical weight distribution for the first time, they generated significantly more compensatory torque at lift onset after observing an erroneous lift (mean = 114.66 ± 8.46 Nmm) compared to having no observation (mean = 17.38 ± 2.77 Nmm) (right side of Figure 5B). However, observing lifting errors did not improve predictive lift planning equally well as haptic feedback, because lift performance of the error observation group was significantly better in the second lift after both error observation (mean = 208.39 ± 9.64 Nmm; p < 0.001) and without observation (mean = 201.57 ± 6.71 Nmm; *p < 0.001*). In contrast, in the skilled observation group, we did not find any evidence that observing skilled lifts on an asymmetrical weight distribution improves predictive lift planning. In their first lift after the unexpected change in weight distribution, participants generated compensatory torque that did not differ significantly between without observation (mean = 34.60 ± 9.89 Nmm), skilled noncollinear (mean = 84.69 ± 10.99 Nmm; *p = 0.47*) and skilled collinear observation conditions (mean = 42.20 ± 8.44 Nmm; *p = 1.00*). Moreover, compensatory torque generation in these first lifts was significantly worse compared to their second lifts (pooled across conditions: mean = 200.38 ± 4.78 Nmm; for each within-condition comparison: *p < 0.001*). In conclusion, these findings indicate that error observation has a stronger impact on predictive planning of compensatory torque on asymmetrical weight distributions than skilled observation. However, these effects are smaller compared to when participants had access to haptic feedback. Noteworthy, these improvements did not lead to reduced object tilt as discussed in the previous paragraph.

##### Digit positioning strategies are similar after both haptic and visual feedback

The grey bars in Figure 5C show that, in the absence of lift observation, both groups positioned their fingertips more noncollinearly in their second lift (skilled observation group: mean = 27.58 ± 2.78 mm; error observation group: mean = 21.75 ± 1.84 mm) compared to their first one (skilled observation group: mean = 0.68 ± 0.45 mm, *p < 0.001*; error observation group: mean = 0.00 ± 0.40 mm; *p = 1.00)* (REPETITION X OBSERVATION_GROUP_: *F*_*(3,228)*_ *= 13.13, p < 0.001)*. As such, both groups preferred to position their fingertips further apart when they had knowledge about the object’s weight distribution based on haptic feedback. The skilled observation group tended to imitate the informed actor’s digit positioning in their first lift after lift observation. They positioned their fingertips noncollinearly (mean = 26.15 ± 1.74 mm) or collinearly (mean = 2.65 ± 1.76 mm) after observing the informed actor using noncollinear or collinear positioning, respectively. Accordingly, in their first lift of the noncollinear condition, the skilled observation group positioned their fingertips significantly further apart compared to their first lifts in the other conditions (*both p < 0.001;* Figure 5C). Furthermore, this noncollinear positioning did not differ significantly from the one the skilled observation group used in their second lift of the without observation condition *(p = 1.00)*. This suggests that observing skilled lifts with noncollinear digit positioning enabled participants to rely on a similar digit positioning strategy they would use when having haptic feedback. The error observation group also positioned their fingertips more noncollinearly after observing a lifting error (mean = 19.45 ± 2.23 mm) compared to having no observation (*p < 1.00*). Moreover, this digit positioning strategy was not significantly different to the one they relied upon when having haptic feedback (second lift of the without observation condition; *p = 1.00*; Figure 4C). By and large, these findings indicate that the error observation group’s digit positioning strategy did not differ significantly after having haptic or visual feedback about the object’s weight distribution. To end, our findings provide no evidence that the error observation group, after observing a lifting error (mean = 19.45 ± 2.23 mm), positioned their fingertips differently than the skilled observation group after observing a noncollinear lift (mean = 26.15 ± 1.74 mm) *(p = 1.00)*.

### 3.2 Center of mass change from asymmetrical to symmetrical

#### 3.2.1 Actor’s lifting performance

Although we were initially interested in whether lift observation can improve predictive object lifting on objects with an asymmetrical weight distribution, we included the center of mass change from asymmetrical to symmetrical for completeness. Here, we included the last (i.e., third) lift on the ‘old’ asymmetrical and the first lift on the ‘new’ symmetrical weight distribution. As the actors’ data was intended to validate our experimental set-up, and results were similar to those in ‘the center of mass change from symmetrical to asymmetrical’, we will only discuss object tilt in detail. All data of the informed and naïve actors can be found in Table 1.

**Table 1.** The actors’ lifting performance for old asymmetrical to new symmetrical. Values represent the lift performance of the informed actor (skilled observation group) and naïve actors (error observation group) in their first and second lift on the new symmetrical weight distribution. Values are presented as mean ± SEM. No statistics are shown in this table.

|  | Informed actor |  |  |  | Naïve actors |  |
| --- | --- | --- | --- | --- | --- | --- |
|  | Old asymmetrical |  | New symmetrical |  | Old asymmetrical | New symmetrical |
|  | Informed<br>noncollinear | Informed<br>collinear | Informed<br>noncollinear | Informed<br>collinear | Blind lift | Blind lift |
| Object roll (°) | 0.56 ± 0.44 | 0.84 ± 0.73 | -3.73 ± 0.57 | -7.78 ± 0.89 | 3.38 ± 0.76 | -11.13 ± 0.85 |
| Compensatory torque<br>(Nmm) | 216.53 ± 4.26 | 266.43 ± 5.43 | 29.61 ± 7.17 | 111.81 ± 12.08 | 207.23 ± 5.62 | 140.19 ± 7.43 |
| Digit positioning<br>difference (mm) | 43.18 ± 1.59 | 16.25 ± 1.80 | 6.00 ± 1.03 | 6.38 ± 0.97 | 22.48 ± 1.79 | 5.94 ± 0.55 |
| Total grip force (N) | 14.10 ± 0.71 | 22.41 ± 0.84 | 18.95 ± 1.15 | 25.55 ± 1.49 | 21.18 ± 1.40 | 22.92 ± 1.55 |
| Load force difference<br>(N) | -1.26 ± 0.23 | 2.47 ± 0.37 | -0.40 ± 0.33 | 0.85 ± 0.45 | -0.03 ± 0.37 | 1.93 ± 0.21 |

##### Object tilt

Because, we are primarily interested in lifting performance differences between actor groups on both weight distributions, we will discuss these findings in light of the significant effect DISTRIBUTION X GROUP *(F*_*(1,136)*_ *= 21.32, p < 0.001)*. First, when lifting the old asymmetrical weight distribution, our analysis provided no evidence that the informed actor and naïve actors induced object roll differently (*p = 0.66*). In addition, when the center of mass changed to asymmetrical, both actor types induced more object roll when lifting the new symmetrical weight distribution than when lifting the old asymmetrical weight distribution (informed actor: *p < 0.001;* naive actors: *p < 0.001*). Furthermore, it is important to note that these values are negative, not positive. This shows that the actors were still generating compensatory torque for an asymmetrical weight distribution. That is, exerting compensatory torque on the symmetrical weight distribution causes the object to roll towards the opposite direction, thus causing negative object roll. Last, even though the informed actor could not fully minimize object roll, he still performed better than the naïve actors when lifting the (unexpected) asymmetrical weight distribution for the first time (*p < 0.001*).

Importantly, the effect of DISTRIBUTION X OBSERVATION_GROUP_ *(F*_*(1,136)*_ *= 4.34, p = 0.40)* was significant as well. When the informed actor had previously used noncollinear positioning on the asymmetrical weight distribution, he induced significantly less object roll when lifting the symmetrical weight distribution compared to when he had previously used collinear positioning (*p < 0.001*). As such, these findings indicate that when the informed actor had to change his digit positioning for the new symmetrical weight distribution (e.g., Figure 2A: Actor Lift 3 to Actor lift 5) he was better at minimizing object roll. To end, irrespective of whether the informed actor changed his digit positioning between the old asymmetrical and new symmetrical weight distribution, he was better at minimizing object roll than the naïve actors in the respective first lifts on the new symmetrical weight distributions (*both p < 0.001*).

For transparency, there is one compensatory torque finding that should be discussed as well. Both skilled and naive actors generated significantly less compensatory torque in their first lift on the new symmetrical weight distribution compared to their last lift on the old asymmetrical one (see Table 1; for each group: *p < 0.001*) (DISTRIBUTION X GROUP: *F*_*(1,136)*_ *= 29.01, p < 0.001*). However, when the informed actor changed from noncollinear positioning on the old asymmetrical weight distribution to collinear positioning on the new symmetrical one, he generated significantly less compensatory torque (mean = 29.61 ± 7.16 Nmm) compared to himself when not changing his digit positioning (i.e., collinear on old asymmetrical weight distribution and then on new symmetrical one; mean = 111.81 ± 12.08 Nmm; *p < 0.001*). In sum, these findings substantiate the difference in naivety between actor types. However, our findings also show that the informed actor performed significantly better when changing digit positioning between weight distributions.

#### 3.2.2. Participants’ lifting performance

Briefly, we investigated the participants’ lift performance on their first and second lift on the symmetrical weight distributions (after it unexpectedly changed from asymmetrical) to investigate whether lift observation improves predictive planning. Again, the participants’ lifting performance on the symmetrical weight distribution can be found in the bottom row of Figure 5.

##### Observing lifting errors improves predictive minimizing of object tilt

We will discuss our findings in light of the significant effect REPETITION X OBSERVATION_GROUP_ (*F*_*(3,228)*_ *= 3.73, p = 0.012)*. Our results provide no evidence that observing skilled lifts improved the skilled observation group’s predictive lift planning. Specifically, the amount of object roll the skilled observation group induced in their first lifts after lift observation (Figure 5D, first green and blue bar) did not differ significantly compared to when they had no visual or haptic feedback about the object’s weight distribution (Figure 5D, first grey bar; mean = –10.39 ± 0.65 °; *both p > 0.34*). In addition, when the actor changed his digit positioning for the weight distributions (noncollinear on asymmetrical to collinear on symmetrical), the skilled observation group did not induce significantly different object roll (Figure 5D first green bar; mean = –6.34 ± 0.82 °) compared to when they lifted after the actor not changing his digit positioning between weight distributions (collinear on asymmetrical to collinear on symmetrical; Figure 5D first blue bar; mean object roll = –7.78 ± 0.89 °; *p = 1.00*). Last, the skilled observation group performed significantly better in their second lifts compared to their respective first lift (Figure 5D; *each respective p < 0.001*). Accordingly, these findings indicate that only haptic feedback, and not lift observation type (i.e., observing a potential change in digit positioning) or observing skilled lifts in general, improved predictive minimizing of object roll in the skilled observation group.

Importantly, the error observation group minimized object roll significantly better after observing a lifting error (Figure 5D first red bar; mean = –3.86 ± 1.21 °) compared to having no lift observation or haptic feedback about the symmetrical weight distribution (Figure 5D first grey bar; mean = –11.13 ± 0.85 °; *p = 0.004*). Furthermore, our findings provide no evidence that the error observation group performed significantly worse in their first lift after lift observation compared to their second lifts of both the error and without observation conditions (Figure 5D second red bar; mean = –2.42 ± 0.68 °; second grey bar; mean = –1.64 ± 0.66 °; *both p = 1.00)*. As such, these findings indicate that observing lifting errors enabled the error observation group to predictively minimize object roll.

##### Only error observation improves predictive planning of compensatory torque

As indicated in Figure 5E, the error observation group generated significantly less compensatory torque after error observation (Figure 5E; first red bar; mean = 64.62 ± 8.45 Nmm) compared having no observation (Figure 5E; first grey bar; mean = 140.19 ± 7.43 Nmm) when lifting the new unexpected symmetrical weight distribution (p < 0.001) (REPETITION X OBSERVATION_GROUP_: *F*_*(3,228)*_ *= 5.40, p < 0.001)*. However, the error observation grouped still generated more compensatory torque after lift observation than in their second lift of the without observation condition (Figure 5E; second grey bar; mean = 33.39 ± 3.92 Nmm) although this difference was not significant (*p = 0.19)*. In contrast, the skilled observation group was not able to generate significantly less compensatory torque after observing the actor switch from noncollinear positioning on the asymmetrical weight distribution to collinear positioning on the symmetrical one (Figure 5E first green bar; mean = 79.72 ± 6.91 Nmm;) compared to having no observation (Figure 5E first grey bar; mean = 127.52 ± 10.14 Nmm; *p = 0.92*). In addition, when the actor did not change his digit positioning for the weight distributions (Figure 5E first blue bar; mean = 111.81 ± 12.08 Nmm), the skilled observation group did not generate significantly less compensatory torque compared to the without observation condition (*p = 1.00*). Last, the skilled observation group generated significantly less compensatory torque in their second lifts of each condition compared to the respective first lifts *(all p < 0.001)*. These findings indicate that the skilled observation group was not able to generate the appropriate amount of compensatory torque after lift observation.

##### Digit positioning strategies are similar after both haptic and visual feedback

Briefly, both groups placed their fingertip significantly further apart in their first lift on the unexpected symmetrical weight distribution (mean = 4.38 ± 0.47 mm) than in their second one (mean = –2.21 ± 0.54 mm; *p < 0.001*) (REPETITION: *F*_*(1,228)*_ *= 0.91, p = 0.44*). Although the effect REPETITION X GROUP was significant *(F*_*(3,228)*_ *= 4.42, p = 0.042)*, the post-hoc analyses failed to reveal any relevant significant differences between groups *(all p > 0.46)*. In conclusion, these findings indicate that both groups changed their digit positioning from their first to second lift based on haptic feedback. In addition, we found no evidence that the type of observed lifts (skilled or erroneous) altered digit positioning differently when the center of mass changed from asymmetrical to symmetrical.

#### 3.3 correlations between object tilt and compensatory torque at lift onset

Fu et al. (2010) reported high correlations (r > 0.89) between compensatory torque at lift onset and peak object roll in their study. To validate our findings, we calculated the correlations between peak object roll and compensatory torque at lift onset as well. For the center of mass change from symmetrical to asymmetrical, our analysis revealed a significant Pearson correlation of r = –0.67 (*p < 0.001*) indicating that the less object tilt was present during lifting the more compensatory torque was generated at lift onset. This is plausible considering that participants needed to offset the external torque induced by the asymmetrical weight distribution. For the center of mass change from asymmetrical to symmetrical, we found a significant r = –0.71 *(p < 0.001)* indicating that the more compensatory torque participants generated, the less (or ‘more negative’) object tilt was induced. Again, these findings are plausible considering that the object’s symmetrical weight distribution does not induce any external torque, thus generating compensatory torque causes the object to tilt towards the opposite side.

## Discussion

Previous studies (Meulenbroek et al., 2007; Reichelt et al., 2013; Buckingham et al., 2014; Rens and Davare, 2019) have substantiated that lift observation can mediate critical information about an object’s weight. It is important to note that these studies only investigated changes in object weight and relied on constrained digit positioning. However, when the weight distribution varies and digit positioning is unconstrained, participants can rely on many valid digit position-force coordination patterns to minimize object roll (induced by the object’s asymmetrical weight distribution). Considering end-state comfort (Cohen & Rosenbaum, 2004) and the importance of minimizing object roll in daily life (e.g., when carrying a tray filled with items), we investigated here, whether individuals can also perceive an object’s weight distribution during observed object lifting and, again, use this information to appropriately update their motor command. Participants were required to lift an object with interchangeable center of mass in turns with an actor. The task goal consisted of lifting the object ‘as skillfully as possible’, i.e., minimize object roll. Participants either observed skilled (i.e., minimized object roll) or erroneous lifts (i.e., non-minimized object roll) which could potentially convey critical information about the object’s weight distribution.

Our results indicate that individuals can extract information about an object’s weight distribution from lift observation. Specifically, when participants observed lifting of a novel weight distribution which was erroneous or skilled with a novel digit positioning, they changed their digit positioning strategy. Furthermore, this visual feedback about the object’s weight distribution drove participants to use the same digit positioning strategy as when they had haptic feedback. However, although these changes in digit positioning indicate that participants could perceive the new weight distribution during lift observation, only the error observation group, and not the skilled observation group performed significantly better after lift observation and when not having haptic feedback. Importantly, our results indicate that the error observation group was particularly able to minimize object roll when the object’s weight distribution unexpectedly changed to symmetrical. Furthermore, for both weight distribution changes, the error observation group was significantly better at generating compensatory torque more appropriately than the skilled observation group. In sum, these findings suggest, in line with previous literature (Reichelt et al. 2013; Rens and Davare 2019), that observation of motor errors drives changes in predictive motor control when lifting objects with an asymmetrical weight distribution albeit to only a limited and imperfect extend.

It has been well-established that observing lifting errors mediates object weight-driven predictive lift planning (Meulenbroek et al., 2007; Reichelt et al., 2013). That is, when two individuals have an incorrect expectation of object weight, the second individual will scale his forces more accurately to the actual object weight after observing the first one making a lifting error. Here, in contrast to these studies, lift performance was primarily quantified by digit positioning and the amount of object roll participants generated. We focused on these parameters based on Fu et al (2010). First, they demonstrated that when individuals can freely choose their digit positioning, they place their fingertips further apart (more noncollinearly) when lifting asymmetrical weight distributions. Second, object roll indicates to which extend participants are able to keep the object’s base horizontal during lifting and, thus, comply with task instructions. With respect to these parameters, our results show that the error observation group could perceive the object’s weight distribution during lift observation: When participants lifted a novel weight distribution for the first time, their digit positioning strategy was similar compared to when they had haptic feedback about the actual weight distribution but different compared to when they could not predict the weight distribution (i.e., without lift observation or haptic feedback). Importantly, as the naïve actors in the error observation group could not anticipate the unexpected center of mass change, they did not place their fingertips according to their preferences as, for instance, they positioned their fingertips on the same height for the asymmetrical weight distributions. As such, it important to note that after lift observation, participants did not position their fingertips imitatively but rather by relying on an internally driven strategy. Critically, observation of lifting errors not only drove participants to update their digit positioning strategy, it also enabled them to predictively generate compensatory torque more appropriately and lift an unexpected symmetrical weight distribution skillfully. That is, after observing a lifting error, participants performed better than without observation. Specifically for the change to symmetrical, during their second lifts, when they could rely on haptic feedback, they did not perform significantly better than without this feedback (in their first lifts). To sum up, our findings show, in line with previous studies (Meulenbroek et al., 2007; Reichelt et al., 2013), that individuals are able to perceive intrinsic object properties during observation of lifting errors. Moreover, our findings highlight that observing lifting errors also allows individuals to partially update their motor command for a new weight distribution. That is, after observing lifting errors, participants were able to generate compensatory torque more appropriately after lift observation than without it. However, they were only able to minimize object roll when the weight distribution changed to symmetrical, but not to asymmetrical.

With specific interest to the observation of skilled lifting, Rens and Davare (2019) showed that observing skilled lifts can improve predictive lift planning, albeit in a smaller manner than observing lifting errors. Here, our results suggest that participants who observed skilled lifting imitated the actor’s digit positioning: When the informed actor placed his fingertips on the same height or on different heights, participants positioned their fingertips similarly. As such, it is plausible that the skilled observation group relied on an imitative strategy whereas the error observation group relied on an internally selected motor command. With respect to minimizing object roll, our findings show that the skilled observation group was not able to use lift observation for improving their own lifts. They could not minimize object roll significantly better after lift observation (compared to without observation). In addition, they did improve significantly from their first to second lifts. As such, having access to haptic feedback enabled participants to minimize object roll better, which indicates that it improved the skilled observation group’s lift planning. In addition,

Buckingham et al. (2014) and Rens and Davare (2019) demonstrated that observing lifting errors mediates object weight better than observing skilled ones. Arguably, superiority of observing erroneous lifts for mediating predictive motor planning is likely driven by the typical kinematic discrepancies between erroneous and skilled object lifting (Johansson and Flanagan 2009; Johansson and Westling 1988). Because of these discrepancies, lifting errors have been argued to be more ‘salient’ than skilled ones, that is, more indicative of actual object weight. In line with this notion, our findings suggest that lifting errors indicate the object’s weight distribution, as shown by participants changing their digit positioning strategy after lift observation. However, the skilled observation group also changed their digit positioning after observing skilled lifts with noncollinear digit positioning. Flanagan and Johansson (2003) demonstrated that individuals target their gaze during observation of hand-actions predictively towards the hand rather than responsively. Accordingly, it has been proposed that individuals anticipate movement components during action observation rather than simply monitoring the entire action sequence by itself. Our findings support these hypotheses: both the skilled and error observation groups updated their lifting strategy after observing a change in digit positioning or the object rolling towards its heavy side respectively. Arguably, this discrepancy between expected and observed lifting behavior drove participants to change their digit positioning strategy. In contrast, participants did not change their digit positioning when observing skilled lifts without a change in digit positioning. Presumably, both the unexpected change in digit positioning and object roll could be considered salient with respect to indicating a center of mass change. Noteworthy, in line with previous studies investigating the mediatory effects of action observation on predictive object lifting, our findings indicate that only observing lifting errors mediates predictive lift planning for objects with complex intrinsic properties, such as an asymmetrical weight distribution.

However, it is important to note that our set-up may not allow us to disentangle whether participants inferred the object’s weight distribution based on contextual cues (e.g., digit positioning or object roll) and retrieved a previously established motor command (e.g., from during the practice trials) or whether they truly integrated the intrinsic object properties into their own internal object representation and generated a new motor command each time. Howard et al. (2013) showed that abstract visual cues can lead to formation and retrieval of distinct motor representations. Furthermore, Reichelt et al. (2013) demonstrated that individuals are also able to predictively plan their own lift better after receiving a verbal cue in the absence of action observation. Combined, these studies suggest that individuals are more likely to retrieve their own distinct motor representations, based on contextual information, than to integrate observed actions and generating new motor commands each time. Specifically for our type of object lifting task, Fu et al. (2014) investigated whether participants can minimize object roll (a) with one hand after repeatedly lifting the object with their other hand or (b) when the center of mass is visibly changed after repeatedly lifting the same weight distribution. In both cases they found that participants alter their digit positioning based on the new condition although they are not able to transfer digit force control. In sum, the findings of Fu et al. (2014) further indicate that individuals rather rely on retrieval of previously established motor commands than on inferring the object’s intrinsic properties and shaping a new motor command. In contrast to these findings, Bursztyn & Flanagan (2008) found that digit force control could be transferred when using the other hand or change the center of mass, as long as the hand was also rotated around the object. This suggests that a memory can be updated into a new motor command, but this is context dependent.

The results of our study could also be explained in terms of memory retrieval. Between-group differences in skilled and erroneous performances could then be explained by differences in the clarity of visual cues. That is, larger differences between the expected and actually observed lift might have driven the participants’ ability to retrieve their previously established motor command based on these contextual cues. In addition, this notion could also explain why the participants’ lift performance is suboptimal in their first lifts after lift observation (Figure 5A and 5D). That is, even though contextual cues enable participants to retrieve their previously stored motor command, short-term storage and forgetting might have decreased participants’ ability to appropriately retrieve the correct motor command (Howard et al., 2013). However, this notion may not fully explain our findings. Even though both the skilled observation group (observing non collinear digit positioning) and the error observation group (observing lifting errors) had access to salient information regarding the object’s weight distribution, the error observation group was significantly better at improving their predictive lifting planning than the skilled observation group. Accordingly, irrespective of whether participants recalled an established motor command or formed a new one, our results do indicate that observing lifting errors enables observers to plan their own actions better than when observing skilled lifts, thus showing differential effects of observing skilled and erroneous actions on predictive lift planning. To address the issue whether a motor command was retrieved from memory or formed from observed information, future studies could include a ‘non-observation’ condition in which participants receive an auditory or visual cue. This would then allow to investigate how action observation differentially affects motor planning than non-action related contextual cues.

One of the limitations of the present study is that our inverted T-shape object only consisted of three compartments for changing the weight distribution. This is a small improvement to many of the observation studies using a dichotomous approach with two weights (for example see: Rens and Davare 2019; Buckingham and Gribble 2017; Reichelt et al. 2013). But as mentioned before, this may not allow us to truly investigate whether participants infer object-related information from contextual cues or integrate observed movement features into their own motor repertoire. Increasing the number of experimental stimuli and the inclusion of a non-observation condition (e.g., auditory cues) could benefit the research teasing apart the mechanisms underlying predictive lift planning based on action observation. A second limitation is that although the informed actor performed significantly better than the naive actors, his performance was suboptimal as object roll remained present during his lifts. As such, it is likely that the skilled observation group was also able to perceive object roll during lift observation albeit in a smaller amount than the error observation group. Critically, although the informed actor’s performance was imperfect, the large differences between our experimental groups indicate that differences between the observed lifting performance types did matter. Specifically, larger deviations from skilled lifting did not enable the error observation group to better plan their own lifts predictively.

Furthermore, it is important to note that the informed actor’s digit positioning in the noncollinear condition was rather unnatural. For instance, our informed actor placed his fingertips further apart than the participants in the study of Fu et al. (2010), who freely selected their digit positioning. This could have potentially biased the skilled observation group in positioning their fingertips differently as well. We did not find significant differences in digit positioning between the skilled observation group after observing a noncollinear lift and the error observation group after observing an error lift (Figure 5C), even though the difference in digit positioning between groups is relatively large. Non-significance could potentially be driven by the relatively small sample size. Irrespective of the statistical outcome, it is plausible that the unnatural digit positioning of informed actor biased digit positioning in the skilled observation group and subsequently altered their lift performance. Critically, the skilled observation group did not perform better than the error observation group (Figure 5A and 5D). As such, it is unlikely that these unnatural cues regarding digit positioning helped the skilled observation group in any particular way. To end, as mentioned in the methods, we decided on this wide noncollinear position as this was roughly the maximum distance on which the informed actor could position his fingertips while still being able to lift the manipulandum comfortably. Accordingly, this positioning enabled the informed actor to position his fingertips consistently while also providing maximum clarity in the noncollinear condition.

Regarding digit positioning, it is important to note that our naïve actors also placed their fingertips further apart than the participants in the study of Fu et al. (2010). Although these actors were instructed to position their fingertips according to their own preferences, they may have been biased by the experimenter’s instructions prior to the experimental task, which led to this digit positioning. Furthermore, not only the actors but also both participant groups positioned their fingertips further apart compared than those in the study of Fu et al. (2010). Accordingly, all participants may have been biased by observed lifting, whether skilled or erroneous, and the experimenter’s instruction. Considering the operationalized experimental conditions, the specifically designed manipulandum and unnatural digit positionings (compared to previous studies), it is unlikely that our findings are truly representative for real-life settings. Although our study provides novel insights in the effects of observation on predictive planning of lifting actions, future research is necessary to tease apart these mechanisms in real-life settings. In addition, rather than having individuals lift objects in turns, valuable insights may be generated when participants transfer (i.e., hand-to-hand giving) objects to each other. This approach may limit participants in mimicking observed digit positioning, thus creating the necessity of developing a unique and novel digit force-position coordination pattern.

In conclusion, in the present study, participants performed an object lifting task in turn with another individual who lifted either skillfully or erroneously. During the task, they were required to lift an object during which they had to minimize object roll by generating the appropriate amount of compensatory torque. Our results highlight that even though lift observation allows observers to perceive an object’s weight distribution, only observation of lifting errors improve predictive lift planning. As such, our findings extend on previous studies (Reichelt et al., 2013; Rens and Davare, 2019) by showing that lift observation does improve predictive lift planning for objects complex object properties, such as an asymmetrical weight distribution.

## Acknowledgements

VVP is funded by an FWO post-doctoral fellowship (grant: 12X7118N).

---

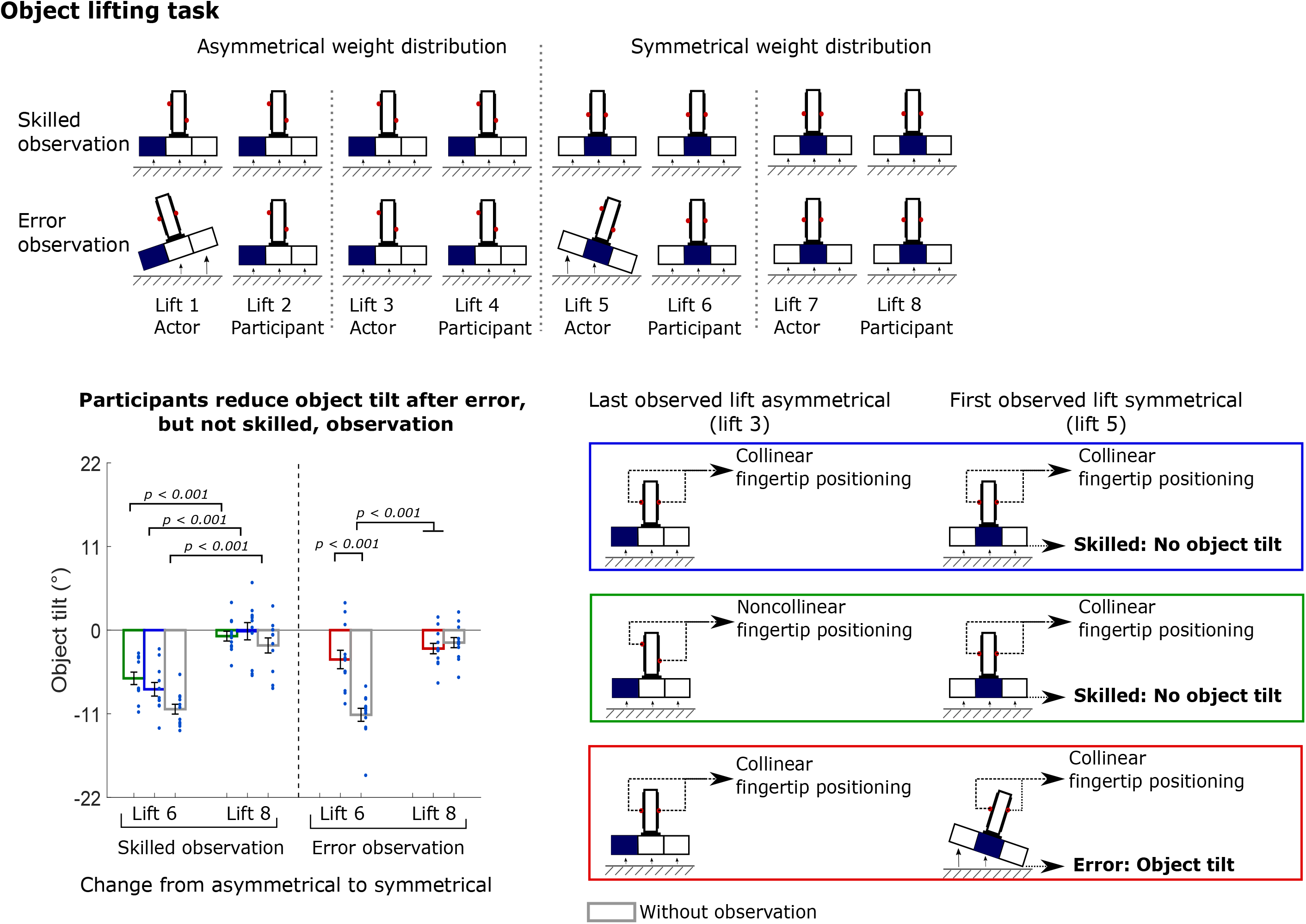

